# Biochemical computation underlying behavioral decision-making

**DOI:** 10.1101/2020.03.14.992057

**Authors:** Stephen C. Thornquist, Maximilian J. Pitsch, Charlotte S. Auth, Michael A. Crickmore

## Abstract

Computations in the brain are broadly assumed to emerge from patterns of fast electrical activity. Challenging this view, we show that a male fly’s decision to persist in mating, even through a potentially lethal threat, hinges on biochemical computations that enable processing over minutes to hours. Each neuron in a recurrent network measuring time into mating contains slightly different internal molecular estimates of elapsed time. Protein Kinase A (PKA) activity contrasts this internal measurement with input from the other neurons to represent evidence that the network’s goal has been achieved. When consensus is reached, PKA pushes the network toward a large-scale and synchronized burst of calcium influx, which we call an eruption. The eruption functions like an action potential at the level of the network, transforming deliberation within the network into an all-or-nothing output, after which the male will no longer sacrifice his life to continue mating. We detail the continuous transformation between interwoven molecular and electrical information over long timescales in this system, showing how biochemical activity, invisible to most large scale recording techniques, is the key computational currency directing a life-or-death decision.

## INTRODUCTION

A fundamental problem in neural computation stems from the need for neurons within a network to communicate without triggering premature responses in downstream circuitry^1–3^. This problem is compounded when the computations are performed over long timescales, as in many decision-making paradigms^4^. In single neurons, the action potential solves an analogous problem by transforming accumulated positive and negative inputs into a transient, all-or-nothing output that is broadcasted to postsynaptic targets. Here we identify a similar mechanism acting at the network level and over timescales ranging from seconds to hours, which we call an eruption.

The Corazonin-expressing (Crz) neurons of the male abdominal ganglion comprise an exceptionally tractable system for investigating neuronal networks and behavioral control. These four neurons drive two simultaneous and crucial events in the lives of the male and his mating partner: i) the transfer of sperm from the male to the female^5^ and ii) a transition out of a period of insurmountably high motivation to continue mating^6^. Both of these events occur six minutes after a mating begins and are under the control of a molecular timer encoded by the slowly-decaying autophosphorylation of the kinase CaMKII^6^. We show that the Crz neurons use cyclic AMP (cAMP) signaling to average evidence about the passage of time across the network and generate an eruption that signals to downstream circuitry only when a consensus is reached. In addition to revealing a new network phenomenon, these results explain the function of CaMKII in the only mechanism for interval timing yet to be described.

## RESULTS

### The Crz neurons form a recurrent network that measures the six-minute time interval

Optogenetic silencing of the Crz neurons (using the light-gated chloride channel GtACR1^7, 8^) that begins before five minutes into a mating results in copulations lasting ∼1-4 hours (long matings), while silencing the neurons any time after seven minutes allows normal, ∼23 minute matings (**Figure 1A**). Long matings result from the failure of the Crz neurons to switch the male out of the state of high motivation that would normally end after ∼6 minutes of mating^6^. This sharp transition is instructed by the kinase activity of CaMKII, which delays the electrical activity of the Crz neurons as it slowly decays over the first ∼6 minutes of mating ^6^ (**Figure 1A**).

**Figure 1:**
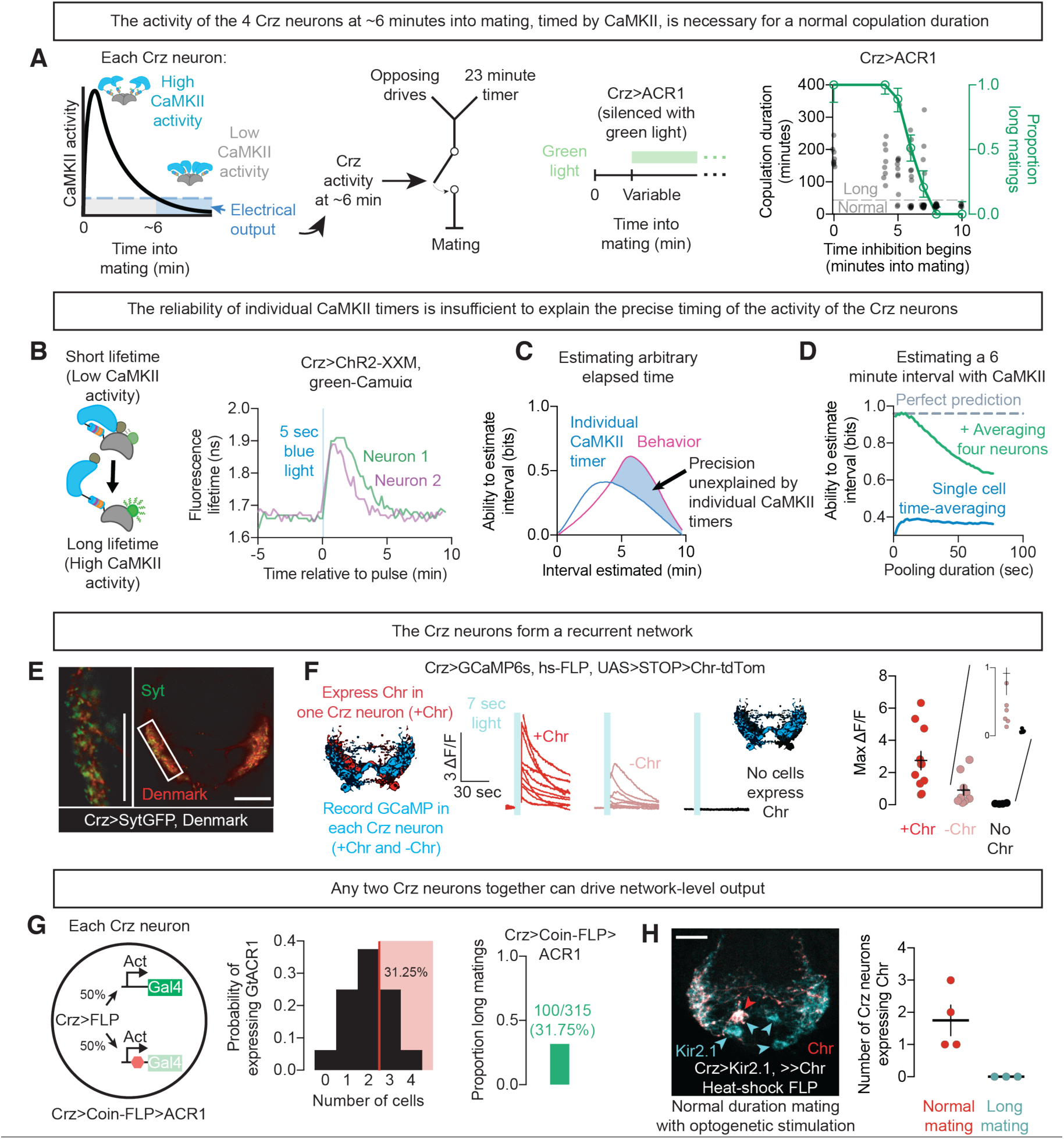
Four Crz neurons form a recurrent network that reports a 6-minute interval. **A)** Far left: The output of the Crz neurons is timed by the declining activity of CaMKII, which takes ∼6 minutes to decay away after the onset of mating^6^. Center left: activity triggers a switch in behavioral state that permits termination in response to both opposing drives and the normal endpoint of the copulation. Center right and far right: Silencing the Crz neurons before 6 minutes into mating extends copulation duration from ∼23 minutes, and often for longer than 1 hour (green), whereas silencing the neurons after 6 minutes has no effect on copulation duration. Individual mating durations are plotted as gray dots; the green curve demarcates the proportion classified as long matings at each time point (see Methods). N values and statistical analyses are listed in **Supplementary Table 2**. Unless otherwise noted, error bars throughout represent 68% credible intervals, selected to approximate SEM. **B)** Simultaneously measured CaMKII timers monitored with green-Camuiα (schematized on left) in two Crz neurons of the same male show similar, but varying, dynamics after optogenetic stimulation using the light-sensitive ion channel ChR2-XXM^9^. **C)** Measuring any individual CaMKII timer is informative about elapsed time, but cannot match the accuracy of the male’s behavior (“ability to estimate interval” is quantified using mutual information, see Methods and **Figure S2**). **D)** Pooling CaMKII activity across time (blue) and across neurons (green) permits a much more reliable estimate of elapsed time, especially if values are compared on the scale of ∼10 seconds. **E)** Synaptic (Synapotagmin::GFP, Syt, green) and dendritic (Denmark, red) markers are closely intermingled in the Crz neurons, suggesting exchange of information. The higher magnification image is taken from the approximate area boxed on the right. **F)** Optogenetic stimulation of individual Crz neurons (dark red traces) excites the other Crz neurons (light red traces), indicating functional recurrence in the Crz network. **G)** Probabilistic silencing (50%, Coin-FLP^10^) of each Crz neuron lengthens copulation in a proportion of males that suggests the requirement of only two active Crz neurons for network function (long matings whenever 3 or 4 neurons are silenced). **H)** Sustained optogenetic stimulation of even a single Crz neuron throughout mating (stochastically selected using a heat-inducible flippase^11^; monitored by tdTomato fusion to Chr; red) is sufficient to recover a normal copulation duration when all Crz neurons are otherwise silenced by expression of Kir2.1 (monitored by EGFP fusion; cyan). This is consistent with the idea that each Crz neuron has access to the output of the network.

Simultaneous measurements of CaMKII activity within multiple Crz neurons of the same fly (using the FRET-FLIM reporter green-Camuiα^12^ after optogenetic stimulation) revealed variability in the duration of sustained CaMKII activity (**Figure 1B**, **Figure S1**), raising the question of how multiple, noisy timers can trigger one-time events like sperm transfer and the abrupt change in motivational state. We quantified the reliability of the CaMKII signal as a timer using an information-theoretic approach^13^ drawing on our previously published green-Camuiα data^6^ (**Figure S2**). We found that measurements of CaMKII from individual Crz neurons cannot account for the reliability seen in the actual fly (**Figure 1C**, **Figure S2B**^6^). Averaging across 4 neurons, however, would allow an almost perfect inference as to whether a 6-minute interval has elapsed, especially if information in the network is exchanged rapidly (**Figure 1D**, **Figure S2C**). This analysis argues that sharing information across the Crz network would, at least theoretically, allow an estimate of elapsed time that reflects the remarkable accuracy observed in the male’s behavior.

One way to share information would be reciprocal excitation, as is suggested by the densely intertwined arborization and intermingled markers for dendritic and synaptic sites of the Crz neurons (**Figure 1E**). We performed a mosaic experiment, expressing the red-light gated cation channel CsChrimson^14^ in a random subset of the Crz neurons while imaging calcium dynamics in all four Crz neurons using GCaMP6s^15^ (**Figure 1F**). In response to red light, neurons expressing CsChrimson showed the expected calcium elevations—but elevated calcium was also always seen in the Crz neurons that did not express CsChrimson (**Figure 1F**). These results do not address whether the connections between the Crz neurons are direct, but they demonstrate functional recurrence and suggest that each of the neurons can receive information about the activity of the others.

If recurrent excitation is important for the network to measure the 6-minute interval, impairing a subset of the neurons should disrupt the system. We generated mosaic animals in which 0,1,2,3, or all 4 of the Crz neurons were silenced in predictable proportions using the recombinase-based tool Coin-FLP^10^, which endows each neuron with a 50% probability of producing Gal4, and therefore expressing GtACR1 for light-gated silencing. If there were a single, special, Crz neuron that triggered the change in motivation, the probability of a long mating would be 50% (the probability of the special neuron expressing GtACR1); if any of the four neurons triggers the switch when its timer expires, long matings should only be seen when all four neurons are silenced (0.5 to the 4^th^ power, or 6.25%). Similarly, if the system requires agreement between *all* neurons, only those flies in which no neurons are silenced would have normal copulation durations (also 6.25%). Instead we found long matings from 31.75% of Coin-FLP-GtACR1 males: 100 long matings out of 315 (**Figure 1G**). This is almost exactly the number predicted if silencing either three (25%) or all four (6.25%) Crz neurons always caused long matings, while silencing two or fewer never caused long matings (**Figure 1G**). This result suggests that no individual Crz neuron is required to produce the event, and that no individual neuron is sufficient to conclude the interval without a partner neuron. The Crz neurons therefore appear to share information not only for the accuracy of the timer, but for its ability to function.

That any two Crz neurons can work together to form a functional timer suggests that all four neurons have the capacity to signal the output. We found support for this idea by expressing CsChrimson in only a subset of Crz neurons while constitutively silencing all four with the leak potassium channel Kir2.1^16^. As long as at least one neuron expressed CsChrimson, sustained optogenetic stimulation could overcome electrical silencing and permit the flies to mate for a normal duration (**Figure 1H**). Any individual Crz neuron can, with sufficiently strong artificial stimulation, change the motivational state of the male, but during mating the network likely relies on recurrent excitation to induce the change. In this model, the output of the Crz network results from the formation of a consensus between at least two of the neurons, achieved by mutual excitation.

### The Crz network uses cAMP to time its output

The Crz network requires ∼60 seconds of electrical activity to throw the motivational switch once the CaMKII timers have run down^6^ (**Figure 2A**). This can be seen by temporarily relieving inhibition after the CaMKII timers have run to completion (10 minutes) and then re-silencing the neurons for the remainder of the mating. If less than 50 seconds of electrical activity is allowed, the motivational switch is rarely thrown (as evidenced by long copulation durations), whereas if more than 75 seconds of voltage dynamics are allowed, the switch is almost always thrown and the mating will terminate ∼18 minutes later (**Figure 2B**, **Supplementary Video 1**). To ask whether 75 seconds reflects the time required for continuous ramping of recurrent electrical activity up to a threshold, we re-imposed optogenetic inhibition between two, independently insufficient, 50-second windows of relieved inhibition. Surprisingly, when one minute of inhibition separated two 50-second windows, males always terminated the mating on time (**Figure 2C**). This demonstrates that continuous activity is not required to throw the motivational switch and suggests the accumulation and storage of a biochemical signal within the network. The accumulated signal often survived even 3 minutes of intervening inhibition, but 10 minutes of inhibition completely erased the effect of the first window (**Figure 2C**). We conclude that depolarization of the Crz neurons drives the accumulation of a long-lasting, likely biochemical, signal that slowly degrades in the absence of electrical activity.

**Figure 2:**
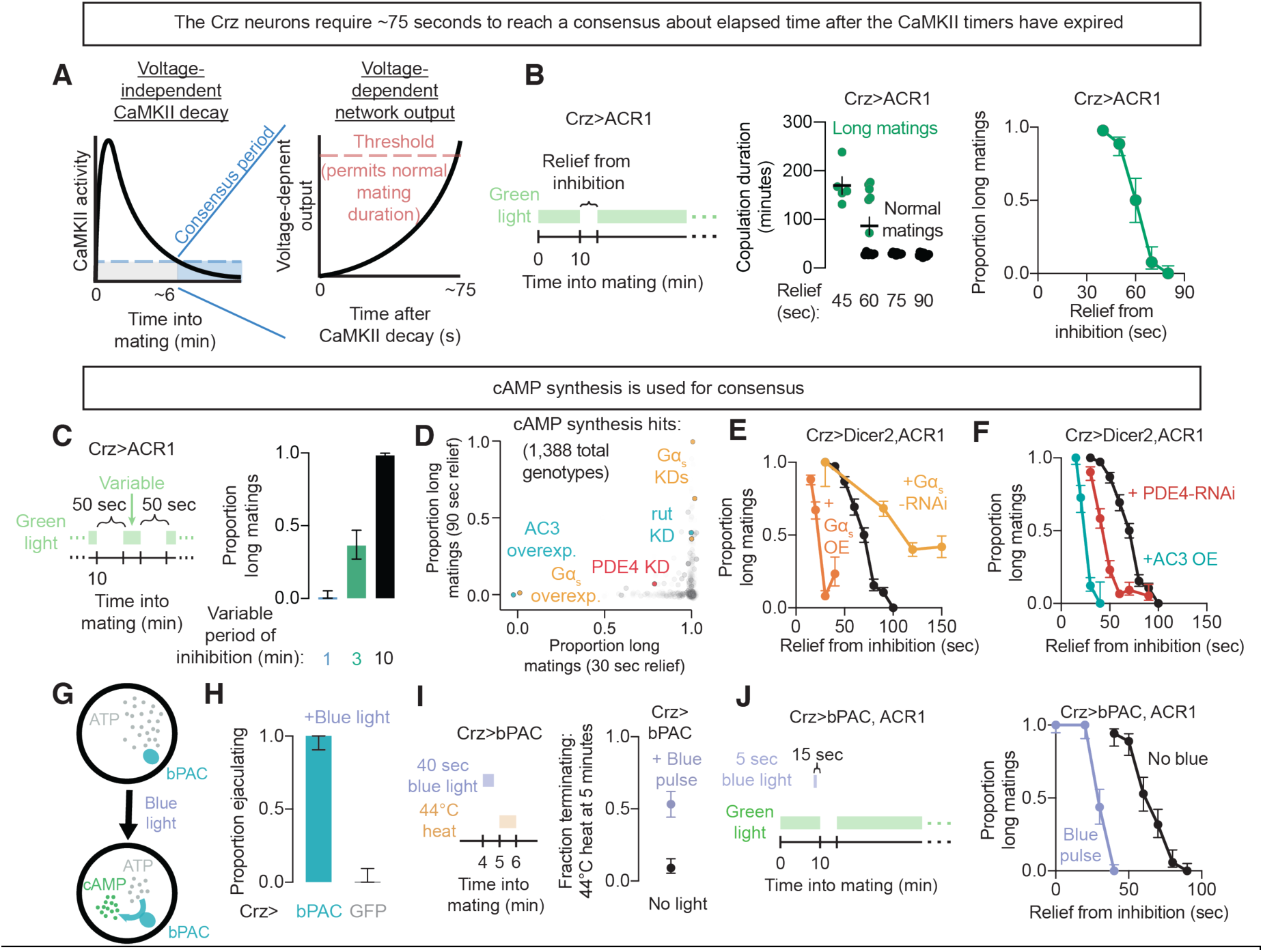
The Crz neurons require cAMP signaling to reach a consensus. **A)** A voltage-dependent signal is required at the end of the CaMKII timer to induce the switch in motivational state. We refer to this as the consensus period due to the requirement for communication between the Crz neurons. **B)** Providing windows of relief from inhibition after the CaMKII timers have expired shows that the Crz neurons require ∼60-75 seconds of electrical activity to induce the motivational switch and cause matings to terminate with normal, ∼23 min durations. If the switch is not thrown, matings last much longer (>50 min). **C)** Reimposing inhibition between independently insufficient 50 sec relaxation windows argues for an accumulating signal that is increased by electrical activity but persists through electrical silencing. The signal can endure 3 (but not 10) minutes of electrical silencing. **D)** A screen through 1,388 genetic manipulations of the Crz neurons (assayed as in **Figure 2B**) uncovered many hits in the cAMP signaling pathway. Colored dots indicate manipulations that are predicted to promote cAMP accumulation and which cause normal matings with only 30 seconds of voltage dynamics (shifting left on the x-axis), or those predicted to prevent cAMP signaling and cause long matings even when 90 seconds of voltage dynamics are allowed (shifting up on the y-axis). rut: rutabaga, an adenylyl cyclase; AC3: adenylyl cyclase 3; PDE4: a phosphodiesterase; KD: RNAi knockdown. Screen data available on request. Further characterization of hits in the screen is provided in **Figure S3**; all screen data is available on request. **E)** Increasing Gα_s_ activity (OE: overexpression), which canonically drives cAMP synthesis, decreases the duration of voltage dynamics required for the Crz neurons to cause the motivational switch, while decreasing Gα_s_ levels increases the time required. **F)** Increasing cAMP synthesis by expression of adenylyl cyclase 3 (AC3 OE) or blocking its degradation by knockdown of phosphodiesterase 4 (PDE4-RNAi), shortens the voltage-dependent period required for the Crz neurons to throw the switch and restore matings to their normal duration. **G)** bPAC, a bacterial photoactivatable adenylyl cyclase, permits light-gated induction of cAMP signaling. **H)** Optogenetic induction of cAMP synthesis in the Crz neurons causes isolated male flies to ejaculate. **I)** bPAC-mediated cAMP synthesis is capable of prematurely inducing the switch in motivational state, as measured by termination in response to potentially lethal heat threats 5 minutes into mating^6, 17^. **J)** Stimulation of bPAC shortens the consensus period, though it does not bypass the necessity for electrical activity to report the output of the network.

To look for a factor that accumulates over time to throw the motivational switch, we screened 1,388 molecular manipulations directed to the Crz neurons (predominantly single gene RNAi knockdowns or expression of mutant *Drosophila* genes, with a focus on calcium-interacting proteins and signaling pathways) for the ability to either shorten or lengthen the necessary duration of the voltage-dependent window (**Figure S3**). Of these, ∼30 showed detectable changes, with many strong hits implicated in the synthesis, stability, and readout of cyclic adenosine-3’,5’-monophosphate (cAMP) (**Figure 2D**). cAMP, a nearly ubiquitous intracellular second messenger in animals, is synthesized from ATP by the action of adenylyl cyclase (which is often activated by the G protein subunit α subtype s (Gα_s_)), and is degraded by phosphodiesterases^18^. Genetic manipulations predicted to impede cAMP synthesis or enhance its degradation lengthened the voltage-dependent time window (**Figures 2D-F**) while those predicted to increase cAMP synthesis or impair its degradation decreased the time requirement (**Figures 2D-F**). cAMP signaling is therefore central to the mechanism that times the motivational switch.

To gain more precise control of cAMP dynamics we employed the photoactivatable adenylyl cyclase bPAC^19^ (**Figure 2G**). Activation of bPAC in the Crz neurons caused ejaculation in isolated males (**Figure 2H**) and was sufficient to hasten the decrease in the male’s motivation during mating^6, 17^ (**Figure 2I**), similar to the effects of thermo- or opto-genetic stimulation^5, 6^. bPAC activation did not have evident behavioral consequences on another population of neurons that are known to drive behavior when stimulated (**Figure S4B**), demonstrating that neuronal activation is not a general consequence of bPAC stimulation. Constant electrical inhibition of the Crz neurons blocked the consequences of bPAC stimulation (**Figure 2J**, 0 sec of relief), showing that cAMP acts through a voltage-dependent process in the Crz neurons, rather than by directly promoting vesicle fusion. Allowing periods of relief from constant inhibition showed that stimulation of bPAC reduced the period required for consensus (**Figure 2J**), suggesting that synthetic cAMP production pools with endogenously generated cAMP to drive the switch in motivation that terminates matings on time.

### cAMP activates PKA to drive voltage-gated calcium currents

A canonical function of cAMP is to activate protein kinase A (PKA)^20^, a tetramer with two regulatory subunits and two catalytic subunits (**Figure 3A**)^21^. cAMP binds the regulatory subunits, causing them to release the otherwise inhibited catalytic domains. RNAi knockdown of the catalytic subunit PKA-C1 lengthened the voltage-dependent time requirement (**Figure 3B**, **Figure S3**). More potently blocking PKA activation with PKA-R*^22^ (a regulatory subunit that cannot be activated by cAMP, and so acts as a dominant negative), lengthened the voltage requirement to the extent that the overall mating duration was increased, often by hours (**Figure 3C**). Like membrane voltage dynamics and the ability to inactivate CaMKII^6^, PKA activity is therefore absolutely required for the Crz neurons to induce the switch in motivation required for appropriately timed mating termination^6^. The dramatic extension of mating duration imposed by inhibiting PKA was overcome by optogenetic stimulation of the Crz neurons (**Figure S5D**), indicating that the neurons remain largely functional, aside from their inability to activate PKA (**Figure S5C**). Inversely, expression of a constitutively active PKA catalytic subunit (PKA-mC*; **Figure 3D**)^22^, but not a wild type or kinase dead version (**Figure S3**), dramatically reduced the duration of the post-CaMKII voltage requirement. These results suggest that PKA reads out cAMP levels to transform the sustained voltage dynamics of the Crz neurons into a discrete output.

**Figure 3:**
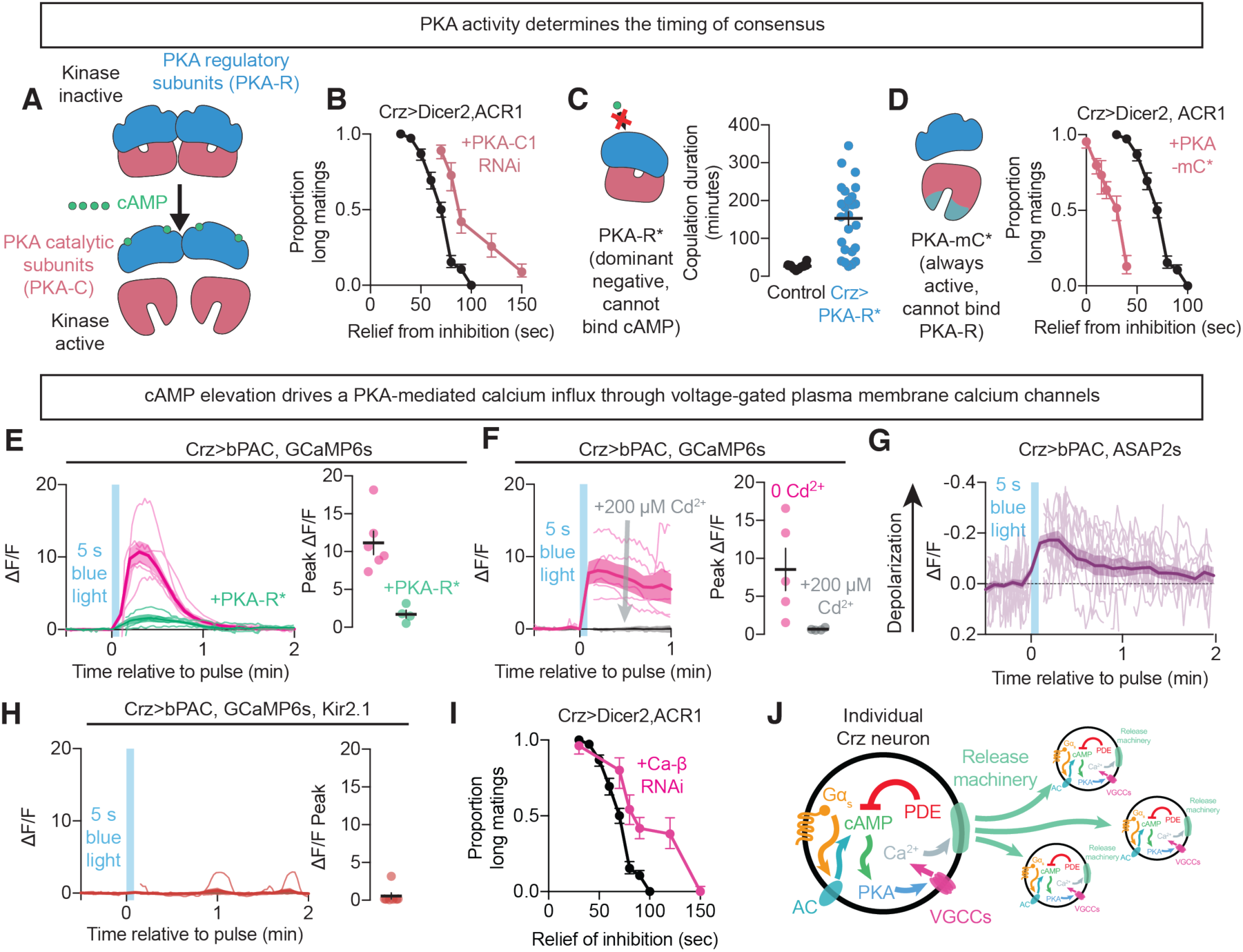
PKA translates cAMP into electrical activity and network output. **A)** PKA forms tetramers of two regulatory subunits and two catalytic subunits. cAMP binds to the regulatory subunits to release the catalytic subunits and permit their action on downstream targets. **B)** Knockdown of the PKA catalytic subunit PKA-C1 extends the duration of the consensus period. **C)** Expression of a cAMP-insensitive PKA regulatory subunit (PKA-R*) prevents the output of the Crz neurons, resulting in matings that last several hours. **D)** Expression of a constitutively active PKA catalytic subunit (PKA-mC*) dramatically shortens the duration of the voltage requirement, though the Crz neurons continue to require electrical activity for the output of the network. **E)** Optogenetic induction of cAMP synthesis using bPAC results in a massive increase in intracellular calcium levels that requires PKA activity (each trace indicates one neuron from separate flies). **F)** Transmembrane calcium influx is required for the PKA/cAMP-mediated increase in intracellular calcium, as the GCaMP6s response is abolished by extracellular application of the Ca^2+^-channel blocker cadmium. **G)** cAMP signaling depolarizes the Crz neurons, as indicated by the voltage sensor ASAP2s. **H)** PKA-induced transmembrane calcium influx requires membrane depolarization, as it is blocked by expression of the leak K^+^ channel Kir2.1. These data together implicate voltage-gated calcium channel signaling in the action of PKA. **I)** Knockdown of the β subunit of the voltage-gated calcium channel lengthens the consensus period of the Crz neurons. **J)** Model illustrating how cAMP signaling drives consensus between the Crz neurons. Gα_s_-signaling activates adenylyl cyclases, resulting in cAMP synthesis. The elevation in cAMP drives PKA activity, which induces calcium influx through voltage-gated calcium channels (VGCCs). Intracellular calcium elevation drives the Crz neurons to signal to each other.

To examine the relationship between cAMP, PKA and neuronal activity, we imaged calcium dynamics in the Crz neurons using GCaMP6s. Sustained stimulation of cAMP synthesis in the Crz neurons using bPAC resulted in extremely large elevations of intracellular calcium that were dependent on PKA activity (**Figure 3E**). This effect required influx through calcium channels in the cell membrane, as the response was eliminated by bath application of the calcium pore-blocker cadmium (**Figure 3F**). The voltage reporter ASAP2s ^23^ showed that bPAC stimulation depolarized the Crz neurons for a comparable duration as calcium influx, indicating that the elevation in calcium is due to a sustained current, not just slow clearance (**Figure 3G**).

Expressing Kir2.1 to hyperpolarize the Crz neurons blocked the change in intracellular calcium evoked by bPAC stimulation (**Figure 3H**), but did not prevent the activation of PKA as measured by the FRET-FLIM tool FLIM-AKAR^24^ that we adapted for use in the fly (**Figure S5E**). The joint requirement of voltage dynamics and membrane calcium channels for cAMP-induced calcium elevation pointed to voltage-gated calcium channels (VGCCs) as mediators of the response to rising cAMP levels. Consistent with this model, knockdown of the *Drosophila* β subunit of the voltage-gated calcium channel complex (Ca-β) lengthened the accumulative window (**Figure 3I**). In mammals, voltage-gated calcium and cation channels are suspected to be modulated by PKA activity to enhance currents^25, 26^, and at least one putative PKA target site on Ca-β is conserved in flies (our observation). However, given the relatively modest effects of Ca-β knockdown, we think it likely that PKA targets multiple calcium conductances to drive the dramatic influx in calcium seen in response to high levels of cAMP signaling.

Multiple non-Crz neuronal populations showed no change in calcium levels in response to bPAC stimulation (**Figure S4**), and the neurons that did respond had much weaker changes in calcium compared to the increase observed in the Crz neurons. The ability of cAMP to excite neurons through the Gα_s_-cAMP-PKA-VGCC pathway (**Figure 3J**) is therefore limited to specific cell types, and the remarkable potency of this pathway in the Crz neurons may underlie the all-or-nothing consequences for behavior and motivational state. Though imaging from the abdominal ganglion during mating remains an experimental obstacle, the correlation between physiological effects in CNS explants and the behavior of the intact animal encouraged us to further explore the network effects of cAMP pathway manipulations.

### cAMP and recurrent dynamics synchronize an eruption of network activity

The behavioral outputs of the Crz network are discrete events (a motivational switch and sperm transfer), while the voltage-dependent period of cAMP accumulation appears continuous. We speculated that this system might employ an accumulate-to-threshold non-linearity, analogous to the action potential in single neurons, so we tested the consequences of driving incremental cAMP accumulation using weaker bPAC stimulation. A 500 ms pulse of blue light produced substantially weaker elevations in intracellular calcium (compared to a 5 second pulse) that returned to baseline within ∼60 seconds (**Figure 4A**). cAMP, in contrast, remained elevated for over 100 seconds 27 (**Figure 4B**), as measured using the cAMP reporter cADDis_GreenDown_ (**Figure S6**, a gift from Vanessa Ruta and Andy Siliciano). cAMP-induced PKA activity measured with FLIM-AKAR lasted even longer (**Figures 4C,D**). Together these data support the hypothesis that cAMP signaling accumulates during the consensus period to build toward the threshold necessary for the output of the Crz neurons.

**Figure 4:**
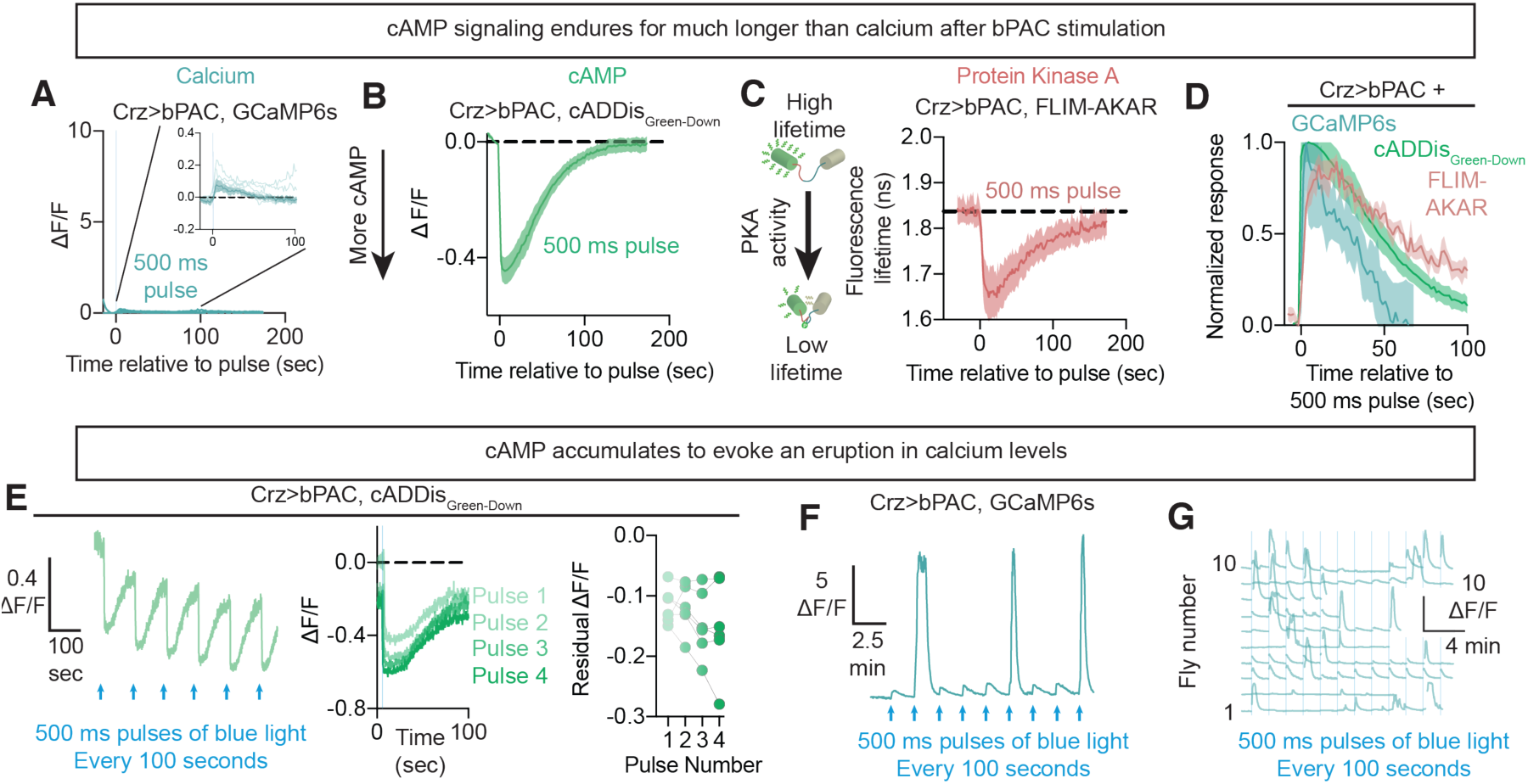
cAMP accumulates within the Crz neurons to drive eruptions of activity. **A-D)** Brief optogenetic induction (a 500 ms pulse) of cAMP synthesis using bPAC elevates cAMP levels (**B**) and PKA activity (**C**) without dramatically changing intracellular calcium (**A**) (inset: zoomed in traces from the larger panel). The resultant elevation of cAMP signaling lasts longer than intracellular calcium (**D**). **E)** Repeated stimulation of cAMP synthesis with bPAC every 100 seconds results in accumulation of intracellular cAMP. Residual cAMP and calcium are computed as the average value during the period 0.25 to 5 seconds preceding the next stimulation (i.e. 95-99.75 seconds after the most recent stimulation for 100 second intervals between each pulse). **F-G)** Accumulation of cAMP signaling eventually results in a massive increase in intracellular calcium levels that we term an eruption. An individual neuron is presented in (**F**), while (**G**) shows the dynamics of several neurons, each from a separate fly.

Consistent with accumulation to a threshold, when 500 ms pulses of bPAC stimulation were applied every 100 seconds (allowing calcium levels to return to baseline) we observed steadily increasing cAMP levels over successive pulses (Figure 4E). Our measurements were often confounded by saturation of the cAMP sensor (**Figure S6**), but accumulation was evident as lingering cAMP just before the next stimulation, when cAMP returns to values within the sensor’s dynamic range (residual cAMP, Figure 4E). Calcium levels responded to this stimulation protocol dramatically differently: roughly every 3^rd^ to 5^th^ pulse triggered what we call an eruption, with massive ΔF/F values approaching the limit of what can be reported by GCaMP6s, and which were sustained for nearly a minute (Figures 4F,G, **Supplementary Video 2**). Eruptions do not result from cell death, as many could be evoked per cell, almost always after several intervening light pulses, and were always of comparable magnitude (Figure 4G**, Figure S7A**). Eruptions were elicited with fewer pulses if the pulses were closer together in time and became very infrequent when separated by several minutes (**Figure S7B**), arguing for an accumulative phenomenon with a slow leak on timescales resembling the behavioral dynamics in our voltage-window experiments. If eruptions were stochastic, on the other hand, we would expect the tendency to erupt to be the same whether pulses were very frequent or very infrequent. To directly examine the relationship between eruptions and cAMP changes, we performed two-color imaging with the red cAMP sensor cADDis_RedUp_ and GCaMP6s. We observed ramping of post-stimulation residual cAMP leading up to the eruption (**Figures S7C,D**). This supports the idea that eruptions result from the accumulation of cAMP itself, likely in association with the accumulation of downstream signaling events.

Thresholds, such as those triggering nuclear reactions and action potentials, can emerge from positive feedback loops if multiple components of a system drive one another until self-sustaining criticality is achieved^28–30^. To look for evidence of such a phenomenon, we first checked whether electrical stimulation of the Crz neurons could increase cAMP levels. Optogenetic excitation of the Crz neurons using CsChrimson 27 decreased fluorescence of cADDis_GreenDown_, reflecting elevated cAMP concentration (**Figure 5A**). These elevations in cAMP levels lasted for tens of seconds, much longer than the calcium response to the same stimulation (**Figure S8**). Repeated bouts of electrical stimulation caused a gradual accumulation of intracellular cAMP (**Figure 5A**). Since the Crz neurons are mutually excitatory, this argues that activity in one Crz neuron elevates cAMP in the others, driving further network activity, forming a positive feedback loop. Though eruptions were most effectively evoked by bPAC activation, eruption-like events could sometimes be driven by direct optogenetic depolarization (**Figure S8**), again consistent with the idea that eruptions are the result of positive feedback between electrical activity and cAMP signaling.

**Figure 5:**
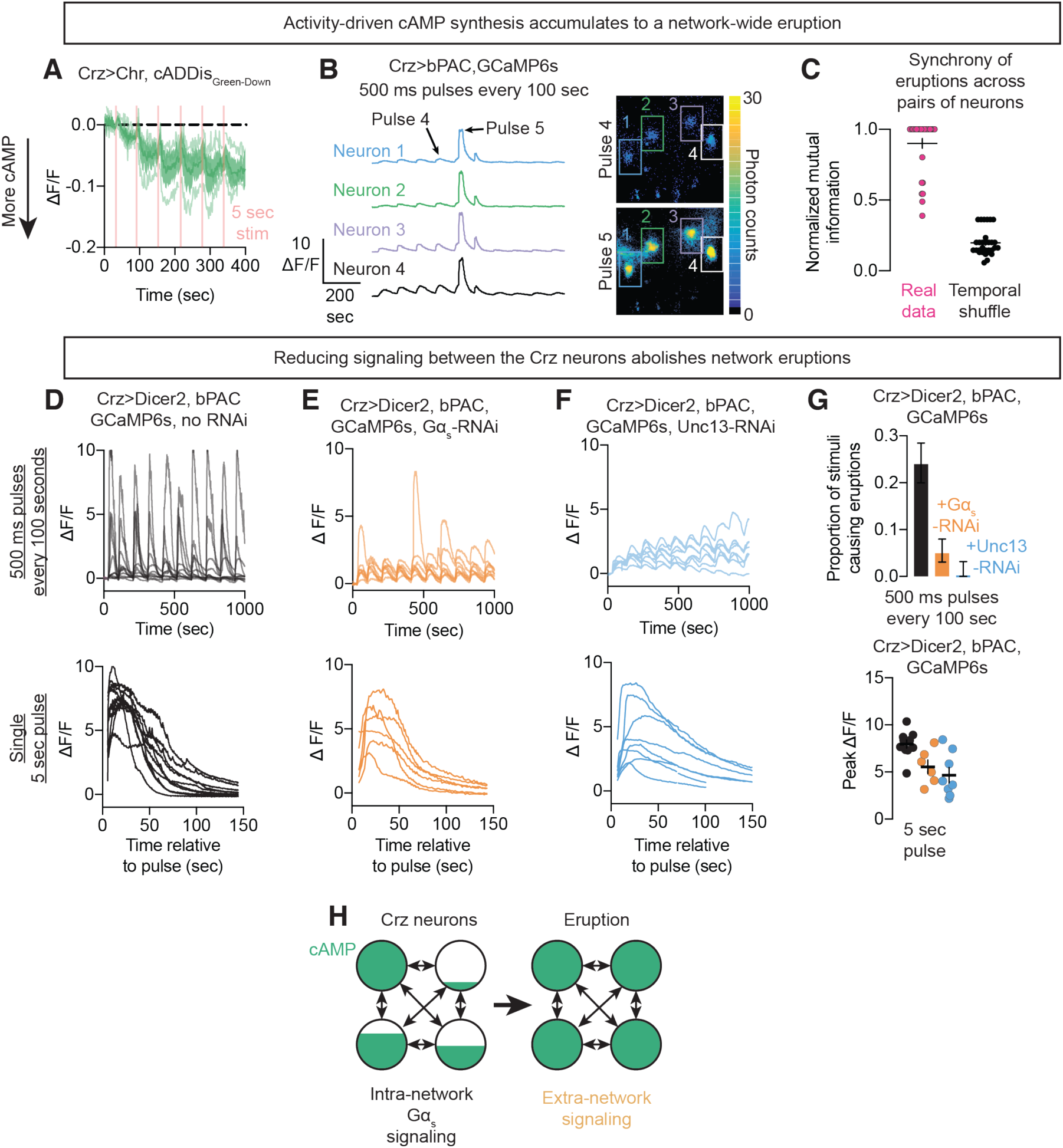
Eruptions are synchronous across the Crz neuron network. **A)** Electrical excitation of the Crz neurons using CsChrimson drives increases in intracellular cAMP levels, as indicated with caDDis_Green-Down_. **B,C)** Eruptions are synchronous across all Crz neurons (for details on the estimation of mutual information, see **Figure S7**). **D-F)** Top row: 500ms pulses of bPAC activation every 100 seconds results in an eruption approximately every fourth pulse on average (**D**). Prevention of vesicle release by knockdown of the active zone protein Unc13 blocks eruptions (**E**).. Knockdown of Gα_s_ within the Crz neurons also prevents eruptions (**F**), demonstrating the requirement of endogenous cAMP synthesis. Bottom row: Prolonged bPAC activation (5 s) can induce eruption-like increases in calcium even when Unc13 and Gα_s_ are knocked-down, demonstrating that the neurons are still largely functional in the absence of these components. **G)** Knockdown of Unc13 and Gα_s_ nearly abolish accumulation eruptions (top row), and reduce the magnitude of induced eruptions from sustained bPAC stimulation (bottom row). **H)** Model for a positive feedback loop between cAMP and electrical activity that concludes with an eruption.

Positive feedback through recurrence suggests synchronization of eruptions across the Crz network, so we were not surprised to find that eruptions in one neuron were a near perfect predictor of eruptions in the other Crz neurons (Figures 5B,C, **Figure S9**, **Supplementary Video 3**). This synchronization likely corresponds to the end of the consensus period after the CaMKII timer and drives the motivational and behavioral responses. To show that eruptions and synchrony were products of recurrent excitation and cAMP synthesis, we stimulated bPAC while depleting genes required for positive feedback from the Crz neurons. RNAi-mediated knockdown of Gα_s_ (Figure 5E) (which would prevent recurrence-mediated cAMP synthesis) or essential active zone machinery (Unc13^31^) (Figure 5F) blocked eruptions that would otherwise be evoked by repeated bPAC stimulation (Figure 5D). Eruption-like events could still be elicited in these configurations by sustained stimulation of bPAC (Figure 5D-G), though often with a smaller response (likely due to impaired recurrence) showing that the neurons remain intact and capable of cAMP-induced excitation. These results together suggest that the eruption is a recurrence-driven network-scale phenomenon.

The Crz neurons are peptidergic, and neuropeptides often require sustained elevations in calcium before release ^32–34^. Our screen (**Figure 2E**, **Figure S3**) was unable to identify any particular transmitter released by the Crz neurons to signal the eruption, but many peptidergic neurons exert their effects through multiple neurotransmitters^35^, raising the possibility of redundancy. Regardless of the identity of the output signal(s), thresholded eruptions likely ensure that the Crz neurons only signal to downstream neurons when the network has reached a consensus, with an unambiguous event that coordinates behavior and motivational state (**Figure 5H**).

### CaMKII delays cAMP-triggered eruptions

The above experiments, together with our earlier demonstration that CaMKII is active for the first several minutes of mating^6^, suggest that CaMKII might time the motivational switch and sperm transfer by delaying cAMP-triggered eruptions. In this model, CaMKII activity serves as a neuron-specific estimation of elapsed time while cAMP levels reflect network-averaged evidence. Neurons with fast CaMKII timers would push the system toward criticality, while neurons whose CaMKII levels remain high delay the output. The network eventually reaches a consensus when the declining CaMKII inhibition can no longer hold back an eruption.

A 90 second voltage window is always sufficient to throw the motivational switch when administered at 10 minutes into mating (or any time after the CaMKII timers have run down). However, even 120 seconds of electrical activity are never sufficient when commenced after only 1 minute of mating, when the CaMKII timers are still highly active (**Figure 6A**). Even the addition of bPAC stimulation had little impact on the ability of voltage windows to allow the motivational switch to be thrown early in mating (**Figure 6B**), suggesting a profound suppression of cAMP signaling by active CaMKII. Artificially elevating CaMKII activity by expression of a constitutively active mutant (CaMKII-T287D) dramatically weakened the behavioral effects of stimulating cAMP synthesis with bPAC, often preventing even tens of seconds of bPAC stimulation from driving the downstream effects (**Figure 6C**). The ability of electrical stimulation to throw the motivational switch, on the other hand, is relatively unimpaired by expression of CaMKII-T287D^6^, further suggesting that CaMKII delays the switch by preventing the accumulation of cAMP signaling, rather than directly preventing electrical activity.

**Figure 6:**
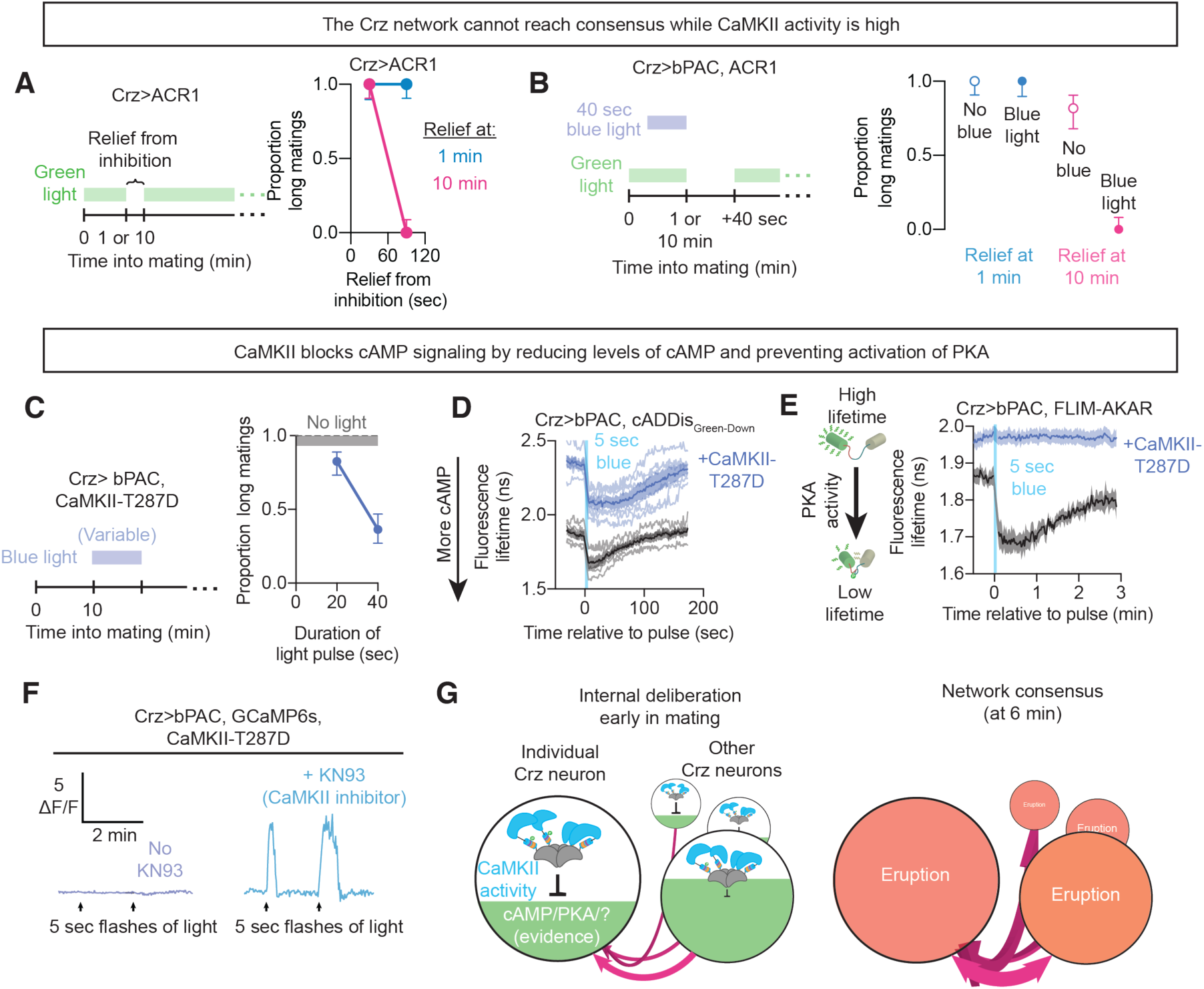
CaMKII delays the eruption by preventing PKA activation. **A)** Windows of relief from optogenetic inhibition cannot trigger the switch in motivational state if provided while CaMKII remains active (1 min into mating). **B)** Optogenetic induction of cAMP synthesis cannot assist in restoring a normal mating duration if delivered while CaMKII is active near the onset of mating. **C)** Expression of constitutively active CaMKII (T287D) dramatically reduces bPAC’s ability to induce the Crz-neuron-mediated motivational switch. **D)** Constitutively active CaMKII reduces cAMP levels to the extent that even 5 second stimulation of bPAC (which would otherwise drive eruption-like events) cannot achieve the normal baseline cAMP levels, as measured with fluorescence lifetime imaging. **E)** Constitutively active CaMKII completely prevents PKA activation in response to strong (5 sec) bPAC stimulation. **F)** Expression of constitutively active CaMKII prevents eruptions evoked by even strong (5 sec) stimulation of bPAC, an effect which can be reversed by application of the CaMKII inhibitor KN93. **G)** Early in mating, each Crz neuron compares two pieces of information to infer elapsed time: its intracellular CaMKII activity, and electrical input from the rest of the network. Network activity drives cAMP accumulation, while CaMKII opposes it. cAMP signaling serves as a common medium for representing evidence that the interval has elapsed. Elevation of cAMP signaling, corresponding to increasing confidence that the interval has elapsed, increases the electrical activity of the neuron, applying pressure to the rest of the network. When recurrent excitation is sufficient to overpower the negating influence of CaMKII activity (i.e. shortly after the 6-minute interval has elapsed), the positive feedback loop between electrical activity and cAMP accumulation results in an eruption, broadcasting the consensus of the Crz neuron network to downstream neurons.

We noticed during earlier experiments that the fluorescence lifetime of cADDis_Green-Down_ changes alongside fluorescence when the sensor binds cAMP (**Figure S6E**). Fluorescence lifetime imaging permits absolute measurements of cAMP levels, instead of the relative changes reported by measuring ΔF/F, since it measures an intrinsic property of individual molecules^36^. We switched to fluorescence lifetime imaging to compare cAMP concentration across flies. The addition of CaMKII-T287D to maintain elevated CaMKII activity dramatically increased the fluorescence lifetime of cADDis-Green-Down (**Figure 6D**), indicating reduced baseline cAMP levels. CaMKII-T287D did not prevent cAMP synthesis by bPAC, but even the maximal level of cAMP achieved after intense stimulation was lower than the baseline levels seen in flies not expressing CaMKII-T287D (**Figure 6D**). This result suggests that active CaMKII prevents cAMP from accumulating to the threshold necessary for PKA activity to generate an eruption. In a striking confirmation of this idea, CaMKII-T287D completely abolished bPAC-mediated activation of PKA (**Figure 6E**), providing a mechanistic explanation for the delayed progression towards an eruption while CaMKII remains active for the first several minutes of mating. At the level of calcium dynamics, we found that imposing high levels of CaMKII activity with the T287D mutation blocked eruptions, even when provoked by strong bPAC stimulation (**Figure 6F**). This block on eruptions was released by addition of the pharmacological CaMKII inhibitor KN93 (**Figure 6F, Figure S10A**). KN93 did not further potentiate eruptions in Crz neurons that expressed a kinase dead phosphomimic mutant CaMKII-K43M-T287D (**Figure S10A**), arguing that the rescue of eruptions is specific to its inhibition of CaMKII activity. These results reveal the mechanism through which the slow decay of CaMKII serves as an interval timer: preventing the accumulation of cAMP signaling from achieving the threshold necessary to report the conclusion of the interval (**Figure 6G**).

### The Crz neurons can accumulate evidence over a wide range of timescales

Evidence accumulation is a broad framework that can describe many systems, unified by the idea of tracking positive and negative inputs over time. Little is understood about the underlying mechanisms, especially on timescales longer than a few seconds. The fact that mating terminates ∼18 minutes after an eruption allows us to use the overall duration of a mating to infer the time at which the network reached a consensus (**Figure 7A**, see Methods). To explore the potential usefulness of this system across modes of evidence accumulation we inhibited the Crz neurons until 10 minutes into mating and then imposed repeated cycles of optogenetic inhibition (negative input) and relief (positive input) (**Figures 7B**). Adjusting the durations of relief and inhibition produced a wide range of copulation lengths, from relatively normal (∼24 minutes) to those lasting many hours. Each cycling inhibition/relief protocol gave a consistently timed output, with more consolidated voltage windows causing earlier eruptions and consequent termination times (**Figure 7C**). Most strikingly, these experiments revealed considerable linearity in the decay of information over time: four seconds of inhibition effectively erased one second of electrical activity across all conditions, even those in which it took over an hour before the network reached consensus (**Figure 7D**).

**Figure 7:**
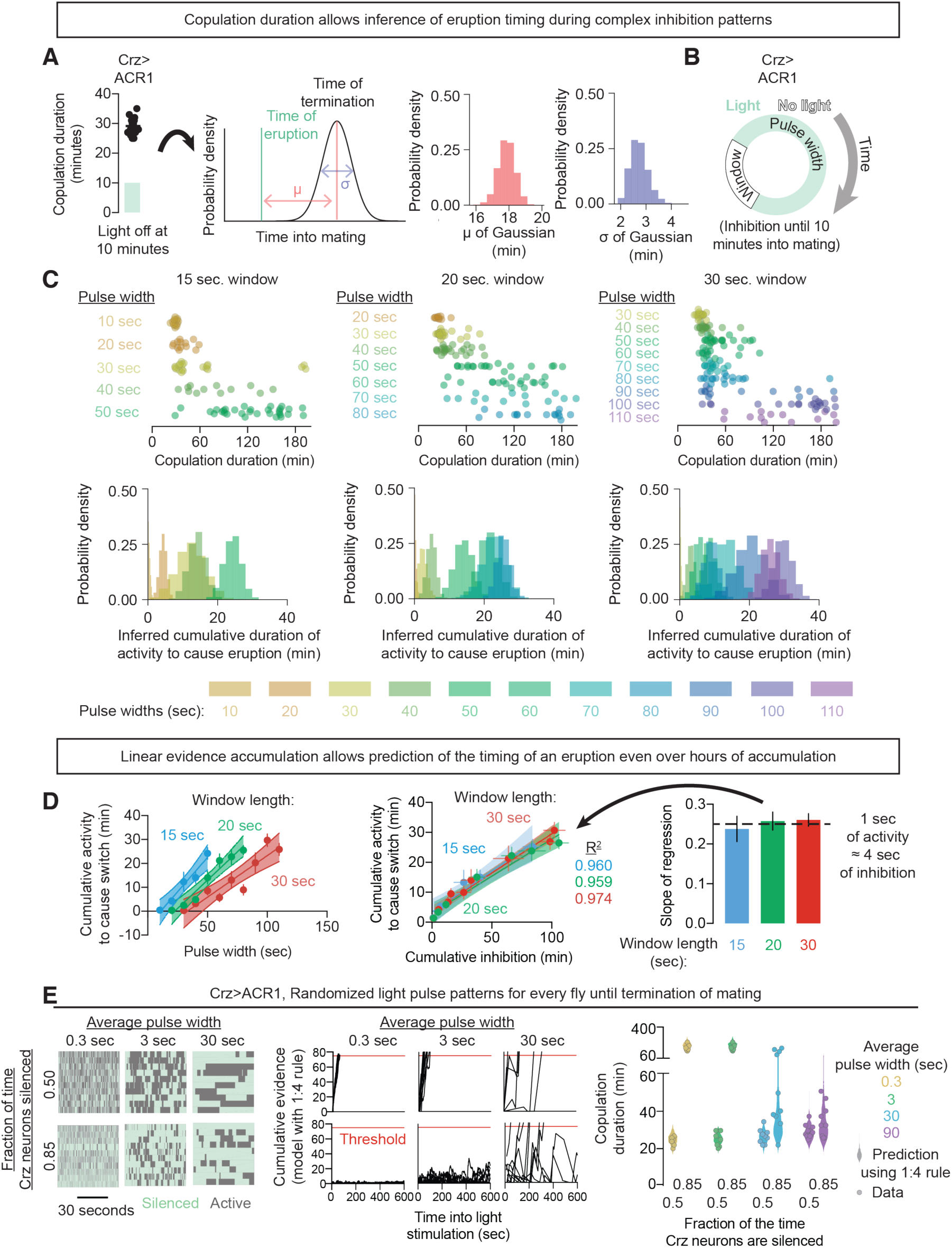
cAMP signaling can track arbitrarily-patterned evidence over a range of timescales. **A)** Delaying the timing of the eruption by optogenetic inhibition permits an estimate of the relationship between an eruption and the cessation of mating. The latency to terminate mating after an eruption was modeled as a Gaussian random variable. The posterior distribution of the Gaussian model’s parameters are presented in red (for the mean) and blue (for the standard deviation). In this instance the Crz neurons were optogenetically inhibited for the first 10 minutes of mating, leading to a presumed eruption at 11 minutes, and a distribution of termination times centered on 29 minutes. Similar ∼18 min delays were seen with various durations of inhibition. **B)** Flies expressing GtACR1 were exposed to 10 minutes of inhibition followed by cycling bouts of inhibition (pulse width) and permitted activity (window length), varying both parameters. **C)** Top row: Individual copulation durations of flies subjected to a fixed window length (indicated at top) and a variety of durations of pulsed inhibition. Longer inhibitory pulses widths always corresponded to a lengthened copulation duration, while increasing the relaxation window length for a fixed pulse width resulted in an earlier termination of mating. Bottom row: Inferred probability distribution for the cumulative electrical activity before an eruption in each condition by using the Gaussian model fit in Figure 7A. Longer copulation durations correspond to a later inferred eruption (see Methods). **D)** Left: For each window length, the cumulative activity required to trigger an eruption increased linearly with pulse width. Middle: Plotting the relationship between total time inhibited and total time active shows a fixed relationship across conditions, with a slope of ∼1/4 – one second of electrical activity negates four seconds of inhibition, regardless of the history of inhibition. This shows that cAMP signaling linearly maintains accumulated evidence, permitting a reliable estimate of total evidence at any point in the consensus process. Right: The constant of proportionality (∼1/4) between activity and inhibition is the same regardless of inhibition paradigm. **E)** Left: optogenetic inhibition was patterned by drawing sequential durations of inhibition and relieved inhibition from an exponential distribution. The mean duration of each pulse of inhibition was specified, as was the average fraction of the time the light was on, but otherwise each pulse was randomized. The randomized phases of inhibition were sustained throughout the mating. Middle: Using only the 1:4 rule derived in Figure 7D, we estimated how long a perfect integrator would need to reach a threshold equivalent to 75 seconds of unimpeded electrical activity (red line). When the fraction of the time the neurons are silenced exceeds 80% (i.e. there is at least 4 times as much inhibition as permitted activity), the integrator struggles to reach the threshold, though the increased variance with increasing pulse width means that for long pulse durations the system can still occasionally reach threshold. Right: Actual copulation duration data (circles) compared to the prediction of the integrator model (violin plots). The model predicts the distribution of copulation durations well, including the possibility of triggering an eruption even with the neurons silenced 85% of the time when the pulse width was long.

If the accumulation and decay of information are linear, the exact pattern of activation and inhibition should not matter, only the predictable time at which the threshold is crossed. To test this idea, we imposed patterned stimuli with programmed averages but otherwise stochastic durations of positive (lights off) and negative (lights on) inputs (**Figure 7e**, **Figure S11**). Using the 1:4 rule (1 second of activity is erased by 4 seconds of inhibition) from the preceding experiments, we could predict the distribution of copulation durations across a wide variety of pulse widths and relative amounts of positive and negative inputs (**Figure 7E**, **Figure S11**). The extreme flexibility of inputs and reliable outputs seen in these experiments suggest that the excitation-cAMP-PKA-VGCC-eruption pathway may be used to track, store, and report information relevant to a variety of behavioral, motivational, and cognitive outputs.

## DISCUSSION

Decisions and behavioral control are thought to arise from long-lasting composite dynamics of neuronal networks^4, 37–40^, such as the seconds-long ramping of firing rates observed in premotor centers^41, 42^. We provide mechanisms for the accumulation and storage of network information over much longer timescales, as well as insight into a thresholding mechanism for reporting the outcome. Here the information is a spatially distributed estimate of elapsed time that emerges from interwoven biochemical and electrical processes. Cell intrinsic evidence is read out within each neuron from the immediate activity of CaMKII, and network-level information is received from the electrical activity of other Crz neurons. The common currency is cAMP signaling, which accumulates intracellularly at the level of cAMP itself, PKA activity, and/or the accumulation of phosphate groups on PKA’s targets. A stark thresholding operation transforms the graded and distributed representation of evidence into a binary decision variable: the presence or absence of a network-wide eruption.

Like action potentials in individual neurons, the system described here rapidly resets after an eruption. In the Crz network, this likely prevents redundant sperm-transfer events. In other systems, the resetting feature may prevent behavioral repetition or inflexibility, while also permitting repeated eruptions if warranted by the strength and constancy of the inputs. Resetting may involve re-activation of CaMKII: elevation of intracellular calcium with strong stimulation of bPAC activates CaMKII (**Figure S10B**), while inhibiting CaMKII with KN93 results in a high frequency of successive eruptions (**Figure S10C**). In mammalian cardiomyocytes, CaMKII is known to phosphorylate and activate phosphodiesterase 4D^43^, which degrades cAMP. There are signs that this mechanism may be at work in the Crz neurons as well: RNAi knockdown of the PDE4 homolog dunce shortens the duration of the voltage requirement (**Figure 2F**), activation of CaMKII dramatically reduces baseline cAMP levels (**Figure 6D**), and a potential CaMKII phosphorylation site on PDE4 seems to be conserved on dunce (our observation). However, it would not be surprising if CaMKII acts at multiple levels to inhibit cAMP signaling, though without strongly affecting baseline membrane voltage^6^. PKA is known to potentiate calcium influx through multiple calcium channels^25, 44, 45^, suggesting a straightforward mechanism for driving the eruption that concludes the timer. But what is the signal that is released by an eruption and delivered to the downstream neurons? It seems likely to be different than the (also unknown) signal used for intra-network recurrent excitation, and could consist of one or more neuropeptides, since they often require sustained excitation for release^32, 46^. Our screens so far have only identified Unc13, a common component of vesicle release machinery for classical neurotransmitters as well as neuropeptides^32, 47^.

Eruption-like mechanisms seem well suited for diverse long-timescale computations. Linearity of evidence accumulation and dissipation allows undistorted evaluation, whether the system is near or far from its threshold. The magnitude of network activation distinguishes the information-gathering phase from the decisive output, allowing evidence to accumulate without triggering the downstream consequences. The accumulated signal persists even in the total absence of electrical input, enabling *activity-silent* memory^48^ that imparts the system with a history dependence that could not be discerned from purely electrophysiological measurements. In some network models, the internal/external signaling problem has been addressed by invoking *output-null*^1, 3^ dimensions of neural activity, but no mechanism has been proposed to control the switch from output-null activity to *output-potent* patterns. Eruptions solve this problem while buffering network-level computations against malfunction of individual components, as demonstrated by our mosaic silencing experiments.

Though we know of no previous description of anything closely resembling an eruption, there are conceptual similarities with several well-studied phenomena in neuroscience. Much of cortex has been hypothesized to operate at near-criticality, giving rise to brief but expansive neural avalanches^49^. Early in cortical development, nascent neuronal networks exhibit spontaneous, network-wide synchronous activation driven by positive feedback that is sustained for tens of seconds^50, 51^. Neurons in primate lateral intraparietal cortex show trial-averaged ramping responses during decision making tasks^39, 52^, but closer inspection shows that individual neurons jump to high activity rates as evidence accumulates^53, 54^. On much longer timescales, neurons in the mammalian suprachiasmatic nucleus (SCN) undergo 10-fold changes in activity depending on time of day^55^. In the SCN, each neuron expresses a cell-intrinsic representation of time of day (encoded by levels of circadian clock proteins), but, as in the Crz neurons, the reliability of behavior is a network output that is much greater than would be expected from its individual oscillators^56^.

We propose to define an eruption as a thresholded jump in recurrent network activity that engages downstream processes that were blind to intra-network accumulation of evidence. This jump in activity can be transient and restricted, as in the four-cell network studied here, and so may have escaped detection in decision making paradigms with less specific labeling capabilities. But we believe eruptions are worth searching for. They might help explain the transformation of continuous brain activity into our discretized actions and experience, and they might refocus our thinking about emergent brain properties. For example, in his influential 1982 description of emergence in neuronal networks, John Hopfield speculated that “the bridge between simple circuits and the complex computational properties of higher nervous systems may be the spontaneous emergence of new computational capabilities from the collective behavior of large numbers of simple processing elements.”^57^. Here we find that profound behavioral and motivational changes instead emerge from the interplay of population dynamics with molecular processing in a remarkably small group of cells. We therefore argue that presuming neurons to be simple processing elements underestimates their computational power. We believe that the computational capacity of individual neurons stems as much from their rich intracellular signaling pathways as from their interconnectedness, especially for computations in the regime of cellular supremacy, where biomolecular computations are proposed to be more efficient than implementations in the classical von Neumann and Turing frameworks^58^. We are confident that studying the molecular/electrical interface in relatively simple systems like the Crz network will provide still more guiding principles for linking the operations of neurons to our thoughts, emotions, percepts, and actions.

### Supplementary Figures

**Figure S1:**
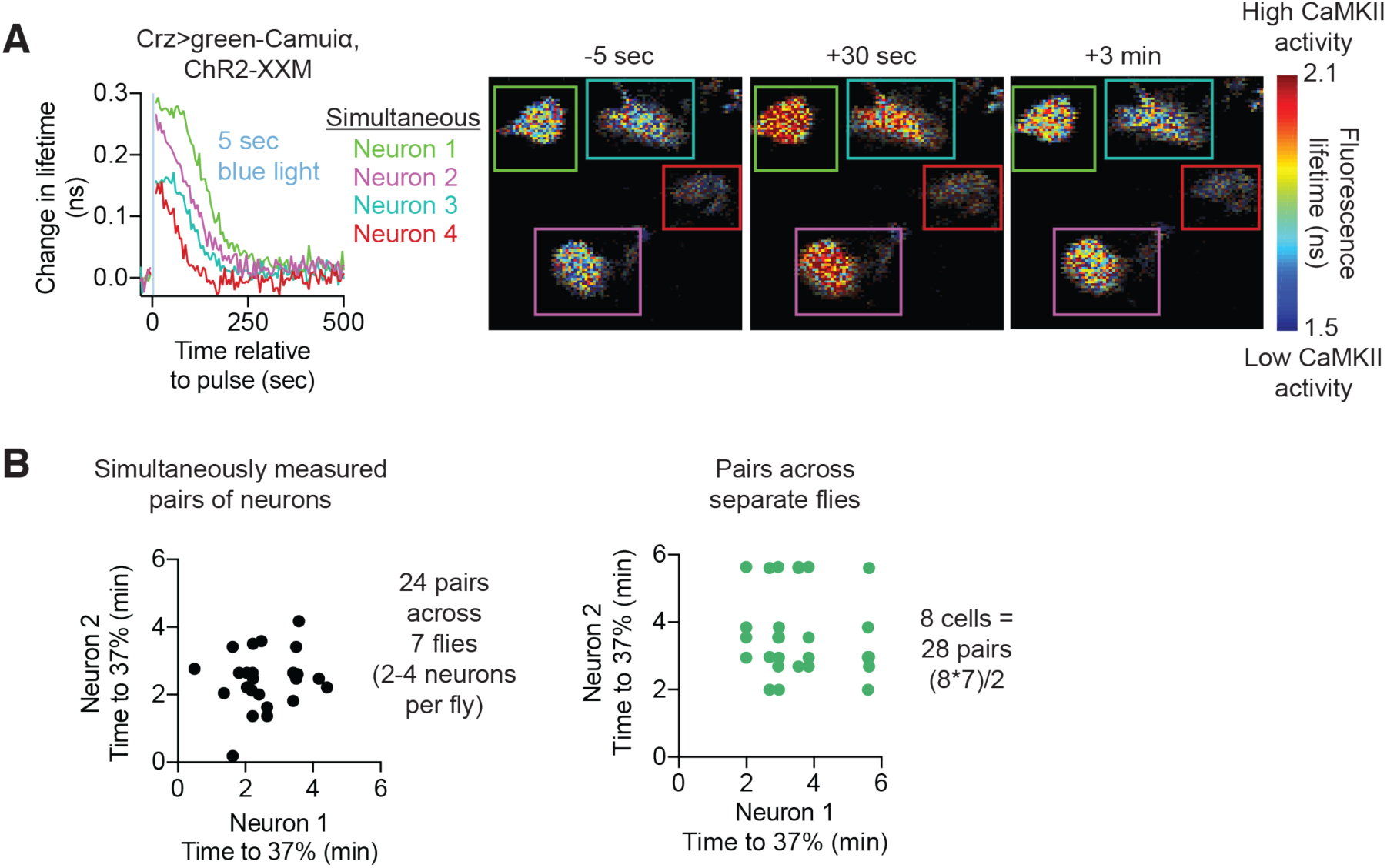
Simultaneously measured CaMKII timers do not report exactly the same interval. **A)** Four simultaneously measured CaMKII timers show high variability relative to the mean decay time of CaMKII. **B)** The variability of simultaneously measured CaMKII timers (left) is comparable to the variability of CaMKII timers across neurons from separate flies (right), arguing that each neuron’s CaMKII timer is independently set.

**Figure S2:**
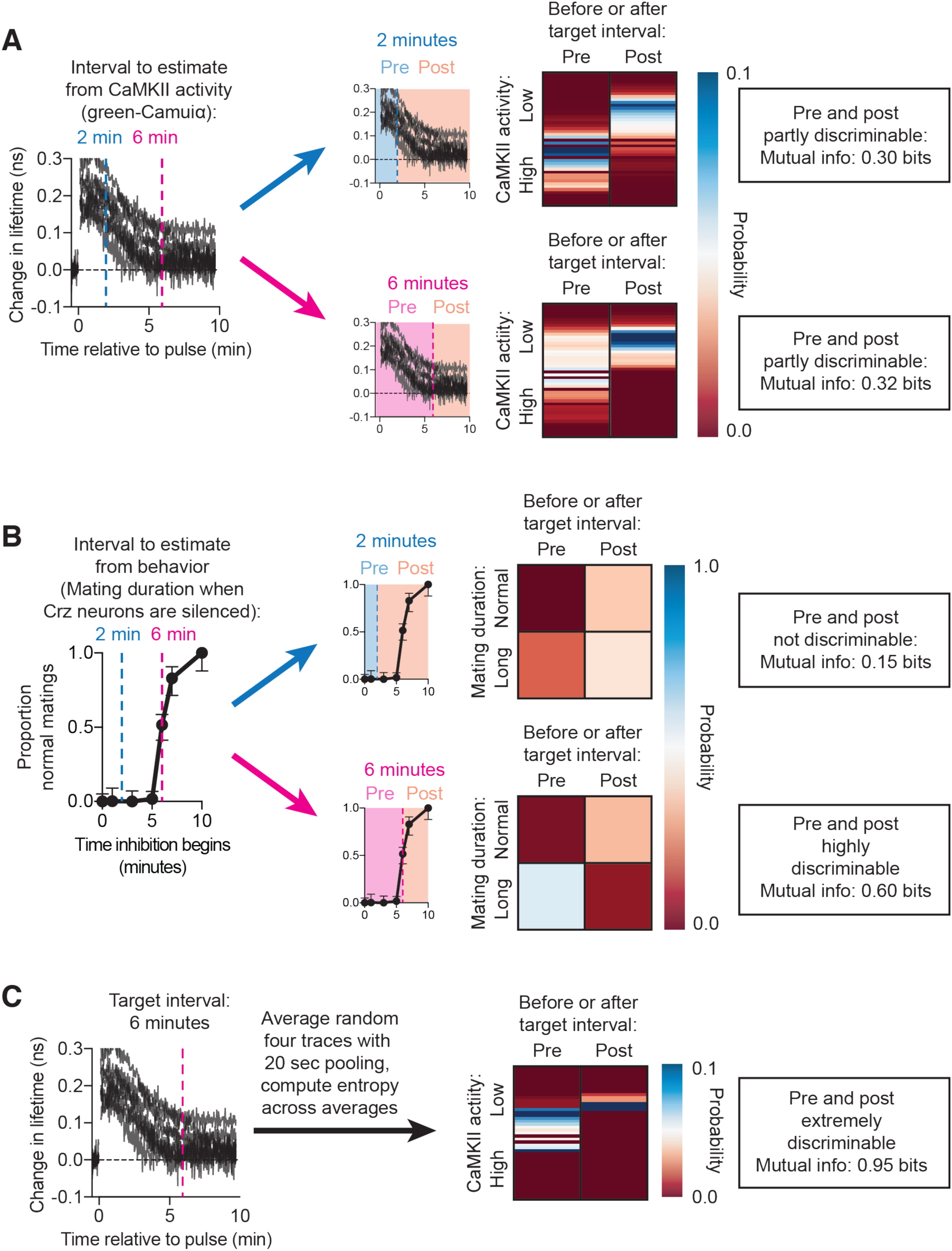
Computation of mutual information between CaMKII timers or behavior and elapsed time. **A)** Eight independently measured CaMKII timers in separate animals after 7 seconds of optogenetic stimulation of the Crz neurons ^6^. To compute the mutual information between CaMKII activity and the passage of a particular interval of time (for example, 2 or 6 minutes), the population of CaMKII responses were split into two epochs: pre and post that interval. The probability density of CaMKII activity in each epoch was computed by binning the green-Camuiα fluorescence lifetime values into 0.02 nanosecond bins. These probability distributions were then used to quantify the entropy of the conditional distribution *p*(Interval has elapsed|CaMKII activity). This was compared to the entropy of the uninformed *p*(Interval has elapsed) in a 10-minute period (i.e. the length of the interval divided by 10 minutes) to compute the mutual information. **B)** To compute the precision of the actual Crz network, we used data in which the Crz neurons were silenced using GtACR1 beginning at various times into mating. If the network has reached consensus by the onset of inhibition, flies will mate for a normal duration^6^. Otherwise, they will mate for hours (long mating). For each target interval, *p*(Interval has elapsed | a fly mates for a long time when the Crz neurons are inhibited) was computed and compared to *p*(Interval has elapsed) as in **Figure S2A**. **C)** As in **B**, but averaging across four neurons at a time.

**Figure S3:**
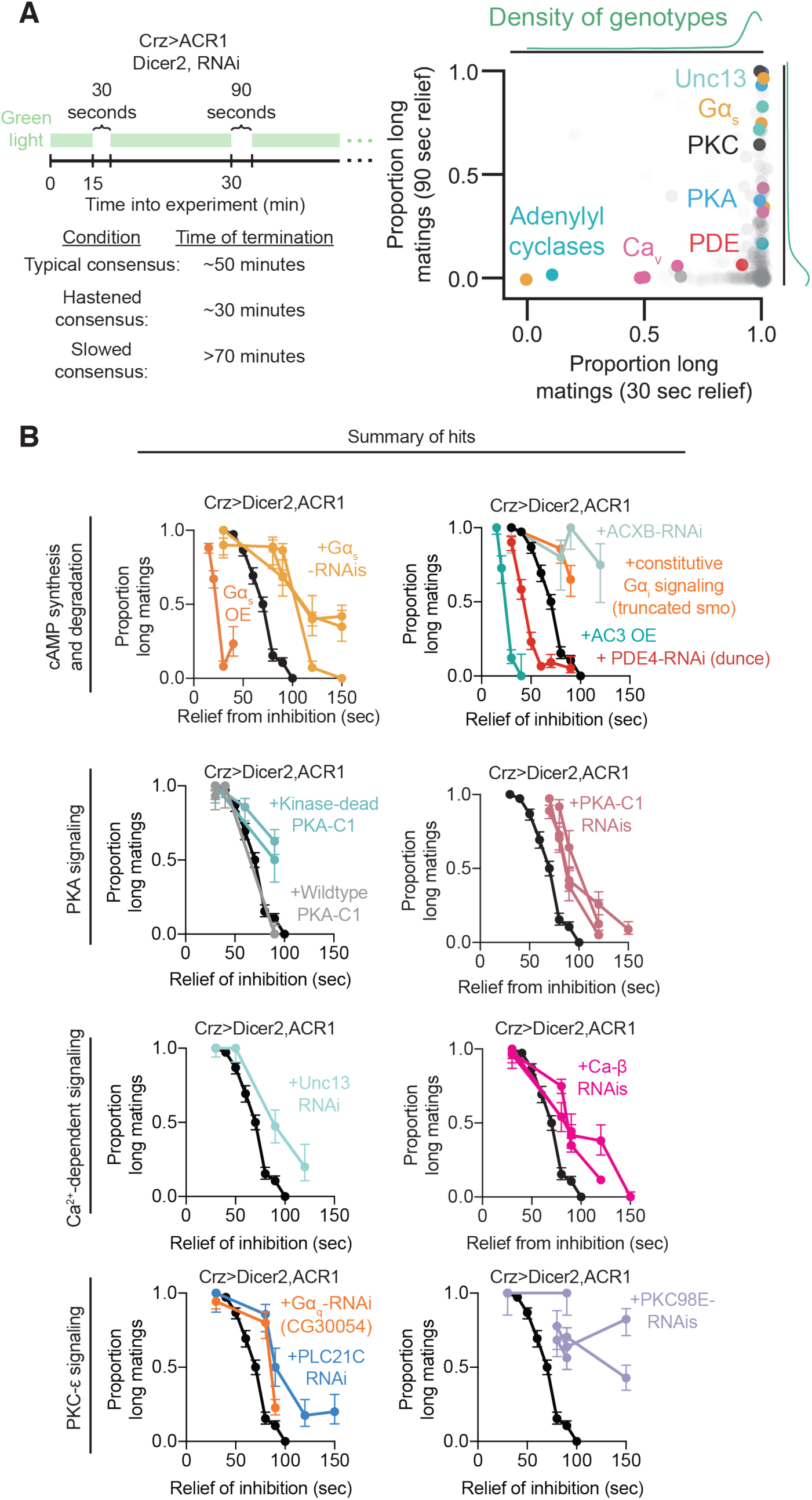
Further details of the screen for the mechanism of the consensus period. **A)** Left: Flies expressing GtACR1 in the Crz neurons, in addition to an RNAi knockdown or overexpression of a single gene, were exposed to two bouts of relieved inhibition: one 30 seconds long, and one 90 seconds long fifteen minutes later. Flies with no genetic modifications other than expression of GtACR1 will persist through the first bout without the Crz neurons reaching consensus, but will respond to the second bout by terminating mating 15-20 minutes later. If the consensus period is shortened by a manipulation, flies will terminate the mating ∼18 minutes after the first period of relieved inhibition. If the consensus period is lengthened, flies will not stop mating even 20 minutes after the second bout. Right: All 1,388 manipulations screened. Most hits were in the canonical cAMP signaling pathway, but we also hit the “novel” Protein Kinase C PKC98E pathway. Opacity of each point increases with sample size (genotypes with more data appear darker in the plot). **B)** Characterization of hits from the screen (with multiple RNAi lines) by plotting the proportion of long matings in response to different durations of relieved inhibition. Each dot represents an independent cohort tested with a different duration period of relieved inhibition.

**Figure S4:**
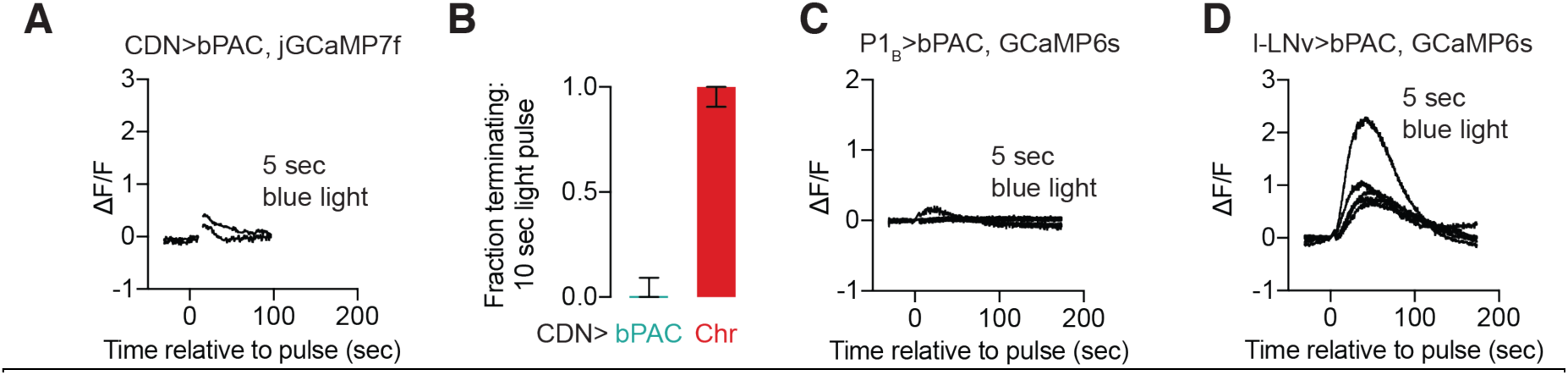
Elevation of cAMP with bPAC does not result in behavioral consequences or eruptions in other neurons. **A)** Stimulation of cAMP synthesis in the CDNs (copulation demotivating neurons (Thornquist and Crickmore, submitted), labeled by the line NP2719-Gal4 in Crickmore and Vosshall ^17^) drives only a modest elevation in intracellular calcium. **B)** Electrical excitation of the CDNs with CsChrimson results in immediate termination of mating, while stimulation with bPAC has no effect. **C)** Stimulation of cAMP synthesis in the courtship-promoting P1 neurons^59, 60^ has no effect on intracellular calcium. **D)** Stimulation of cAMP synthesis in the circadian large lateral ventral neurons (l-LNvs) ^61^ does result in significant rise in calcium, though the increase is much slower and of lower magnitude than in the Crz neurons. This is consistent with the large and rapid increase in calcium in response to bPAC activation in the Crz neurons coming from a threshold nonlinearity between cAMP signaling and calcium. Repeated weak stimulation of the l-LNvs does not show an accumulation-to-threshold or eruption-like effect.

**Figure S5:**
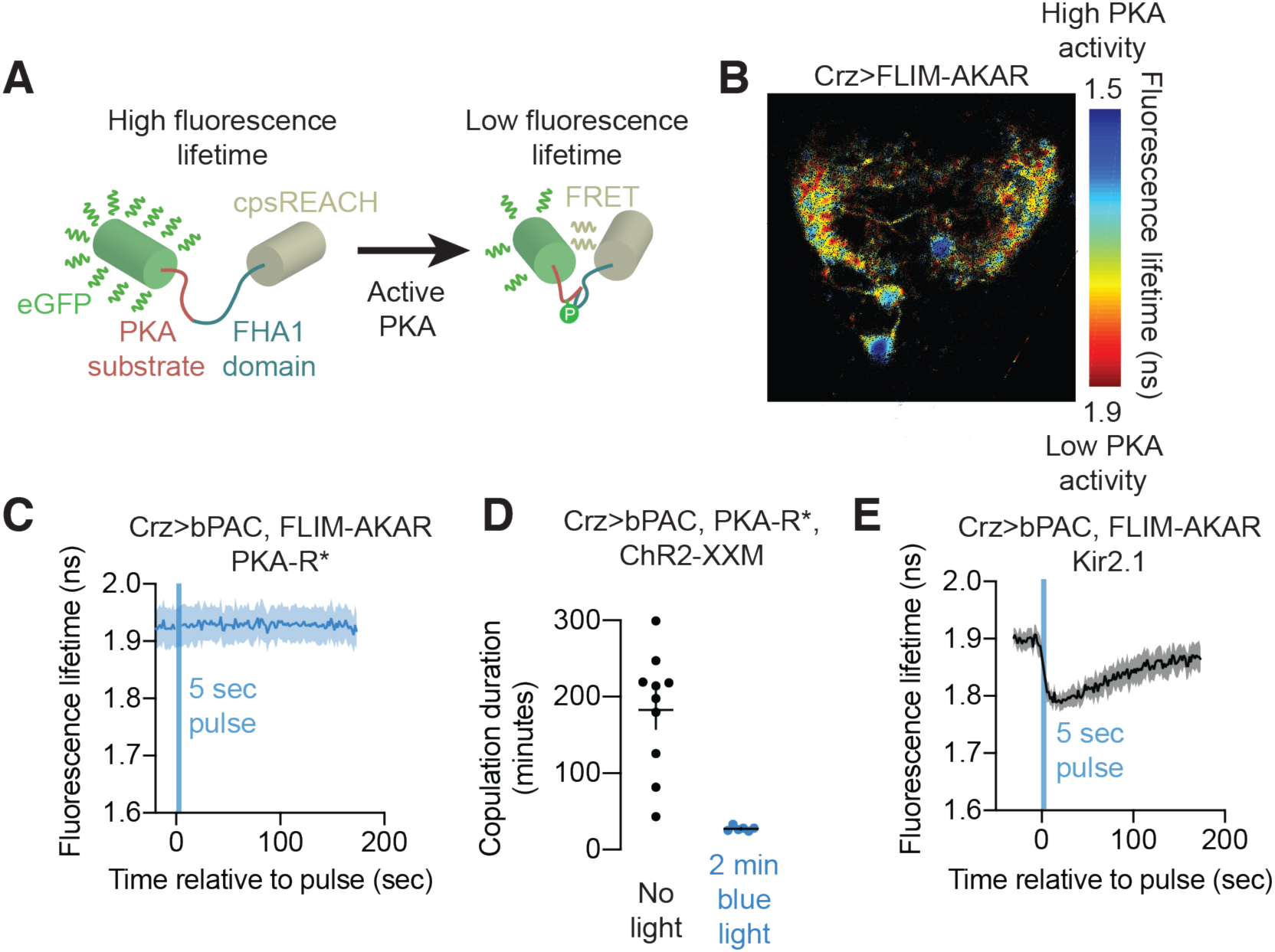
Additional information related to PKA in the Crz neurons. **A)** Phosphorylation of a PKA substrate results in a conformation change of FLIM-AKAR, permitting non-radiative decay of eGFP excitation through FRET ^24^. This is read out as a reduction in fluorescence lifetime, providing a measurement of absolute levels of PKA signaling. **B)** FLIM-AKAR expresses stably throughout the neuron and reveals variation in baseline levels of cAMP within the Crz neurons. Somas show much higher levels of basal PKA activity than the processes, though the dynamics in response to optogenetic induction of cAMP synthesis using bPAC are comparable. **C)** Expression of the dominant negative PKA regulatory subunit PKA-R* completely blocks phosphorylation of FLIM-AKAR after stimulation of cAMP synthesis by bPAC. **D)** Stimulation of the Crz neurons reverts the effects of expression of PKA-R* on copulation duration. **E)** Electrical inhibition of the Crz neurons using Kir2.1 does not prevent cAMP-induced activation of PKA, despite the absence of changes in intracellular calcium (Figure 3H).

**Figure S6:**
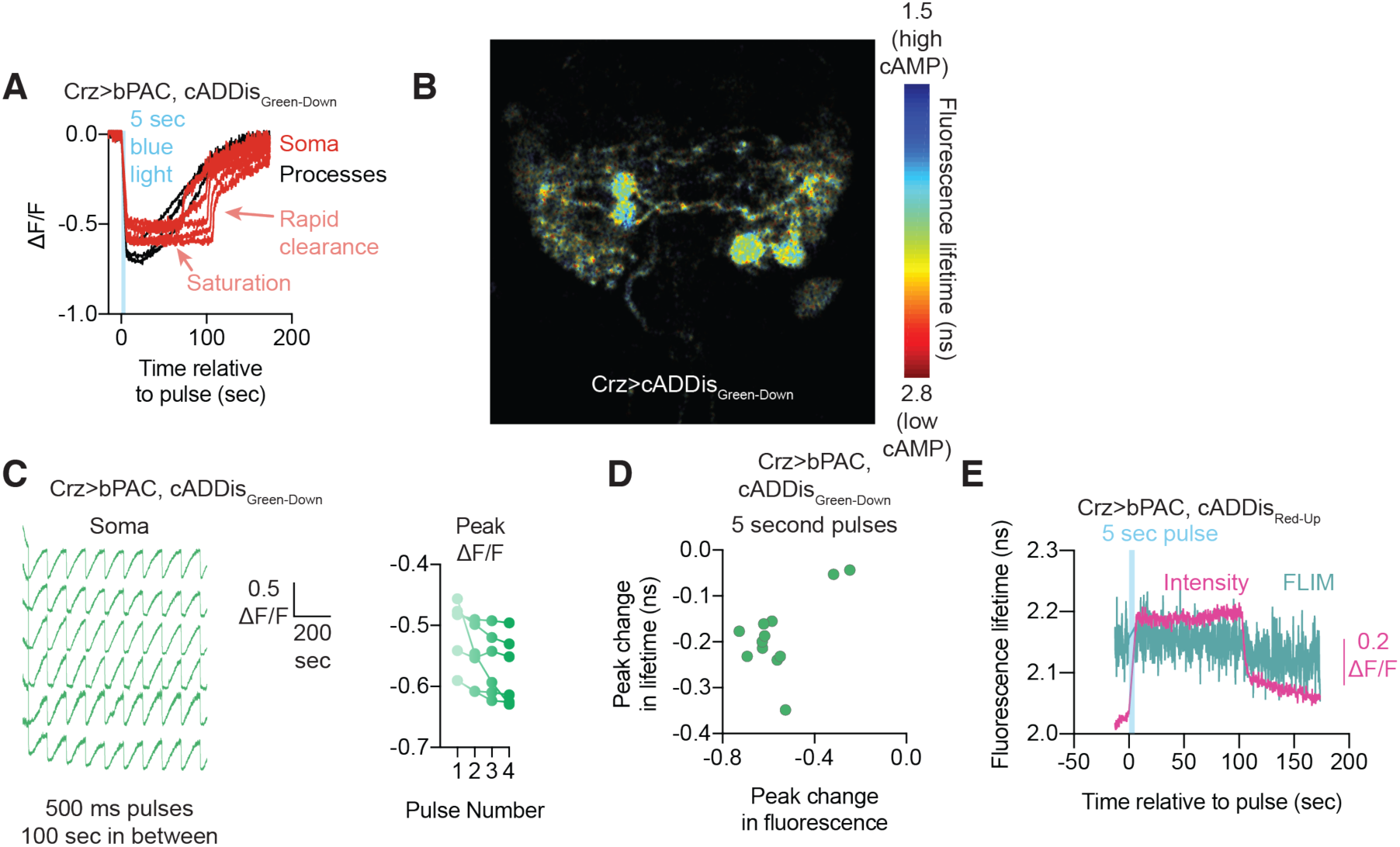
Characterization of the cAMP reporters cADDis_Green-Down_ and cADDis_Red-Up_ in the Crz neurons. **A)** cADDis_Green-Down_ reports large changes in cAMP after strong (5 second) stimulation of bPAC that persists for over 100 seconds. At ∼100 seconds – right at the time that the resultant eruption from optogenetic stimulation begins to end – somatic measurements of cAMP show extremely rapid clearance and return to baseline, suggesting the eruption triggers an active mechanism for cAMP clearance. **B)** Baseline cAMP levels are higher in the soma than in the processes of the Crz neurons, as reported by fluorescence lifetime imaging of cADDis_Green-Down_. **C)** Individual cell traces from data reported in Figure 4E. **D)** cAMP responses in the arbors of the Crz neurons show similar dynamics to the soma, though with less pronounced saturation at the peak of stimulation (likely due to the lower baseline before stimulation).

**Figure S7:**
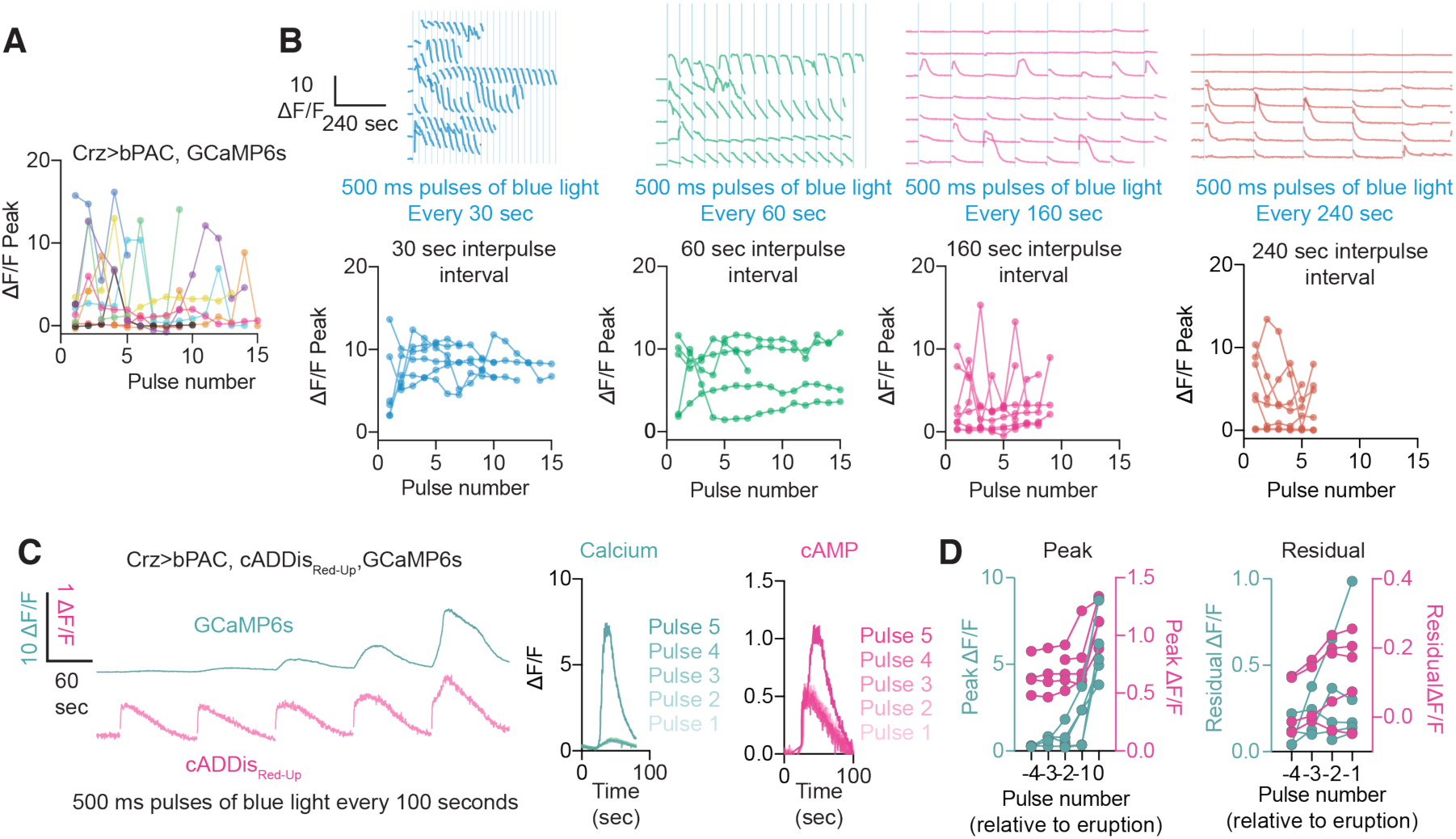
Changing the rate of cAMP accumulation changes the frequency of eruptions. **A)** Responses from individual neurons in Figure 4G are replotted in different colors for comparison of peak heights. **B)** Top: Increasing the frequency of bPAC stimulation eruptions, while further spacing the pulses causes less frequent eruptions. Bottom: Sequential eruptions neither increase nor decrease in magnitude, suggesting that the eruption results from a saturating nonlinearity. **C)** Simultaneous imaging of cAMP and calcium dynamics in an individual Crz neuron shows the relationship between accumulating cAMP and eruptions. **D)** Peak and residual levels of calcium and cAMP during the same experiment across multiple flies

**Figure S8:**
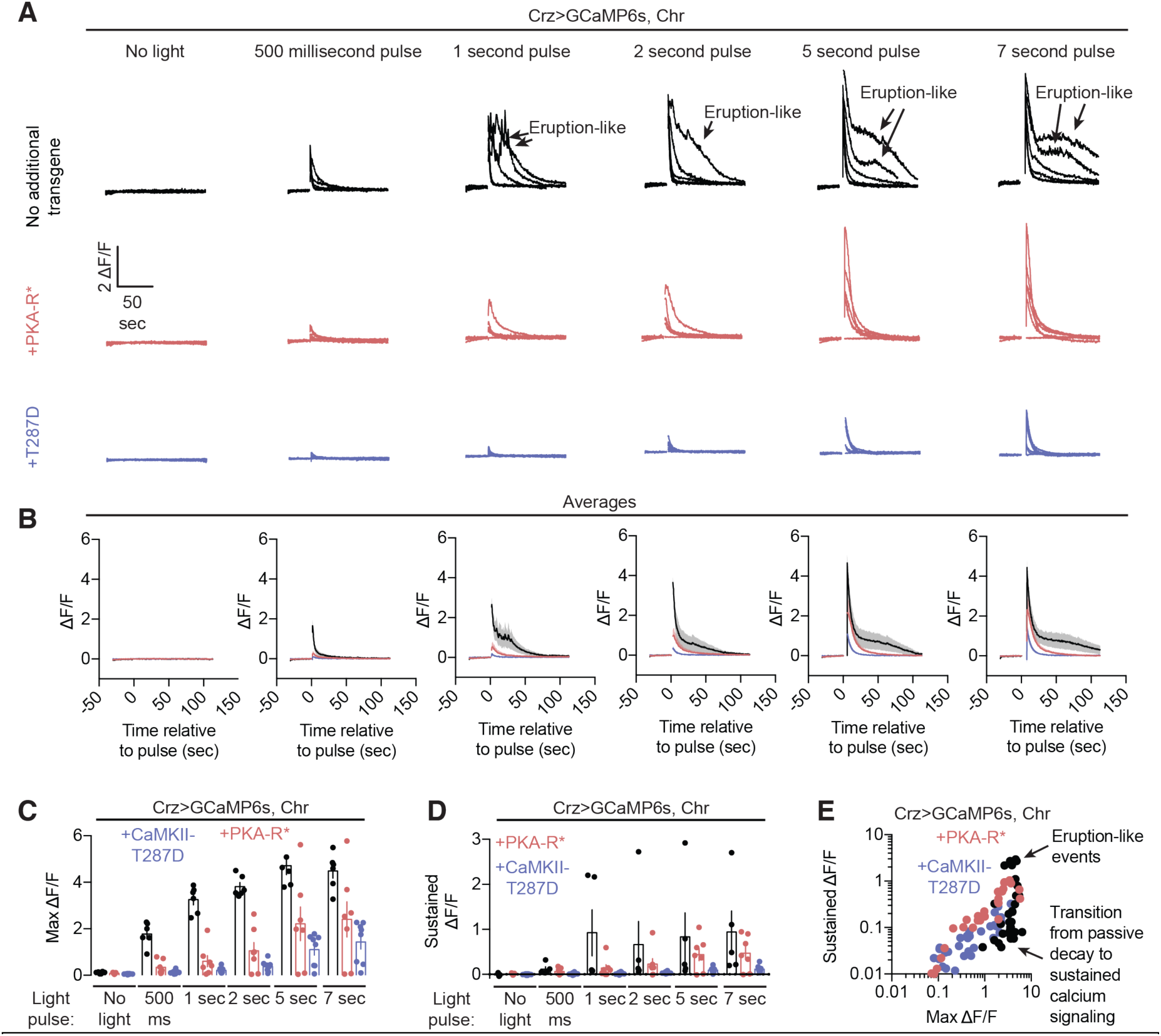
PKA and CaMKII regulate the magnitude of calcium responses to electrical excitation of the Crz neurons. **A)** Individual GCaMP6s responses to optogenetic stimulation of the Crz neurons using CsChrimson. **B)** Average of traces in **Figure S8A**. **C)** Expression of PKA-R* or CaMKII-T287D attenuate the peak elevation of intracellular calcium levels after electrical excitation of the Crz neurons. **D)** Constitutively active CaMKII diminishes the sustained elevation of calcium after stimulation (the average elevation over the period 15-20 seconds after stimulation) ^6^ possibly derived from recurrence, while PKA-R* has a less pronounced effect on sustained activity. **E)** The relationship between the maximal response of the Crz neurons to the sustained response suggests two regimes: one in which the sustained activity is highly sensitive to changes in maximal excitation (when the maximal activity is high), and one in which it is relatively insensitive (when the maximal activity is low). This is suggestive of a transition into recruiting recurrent excitation when the Crz neurons are driven with a sufficiently strong impulse. Below the threshold, recurrence is not recruited and the residual calcium presumably results exclusively from the decay of the initial impulse. Arrow points to the abrupt kink in the relationship between peak and sustained fluorescence changes, indicating when sustained calcium begins to grow rapidly with small increases in stimulation strength.

**Figure S9:**
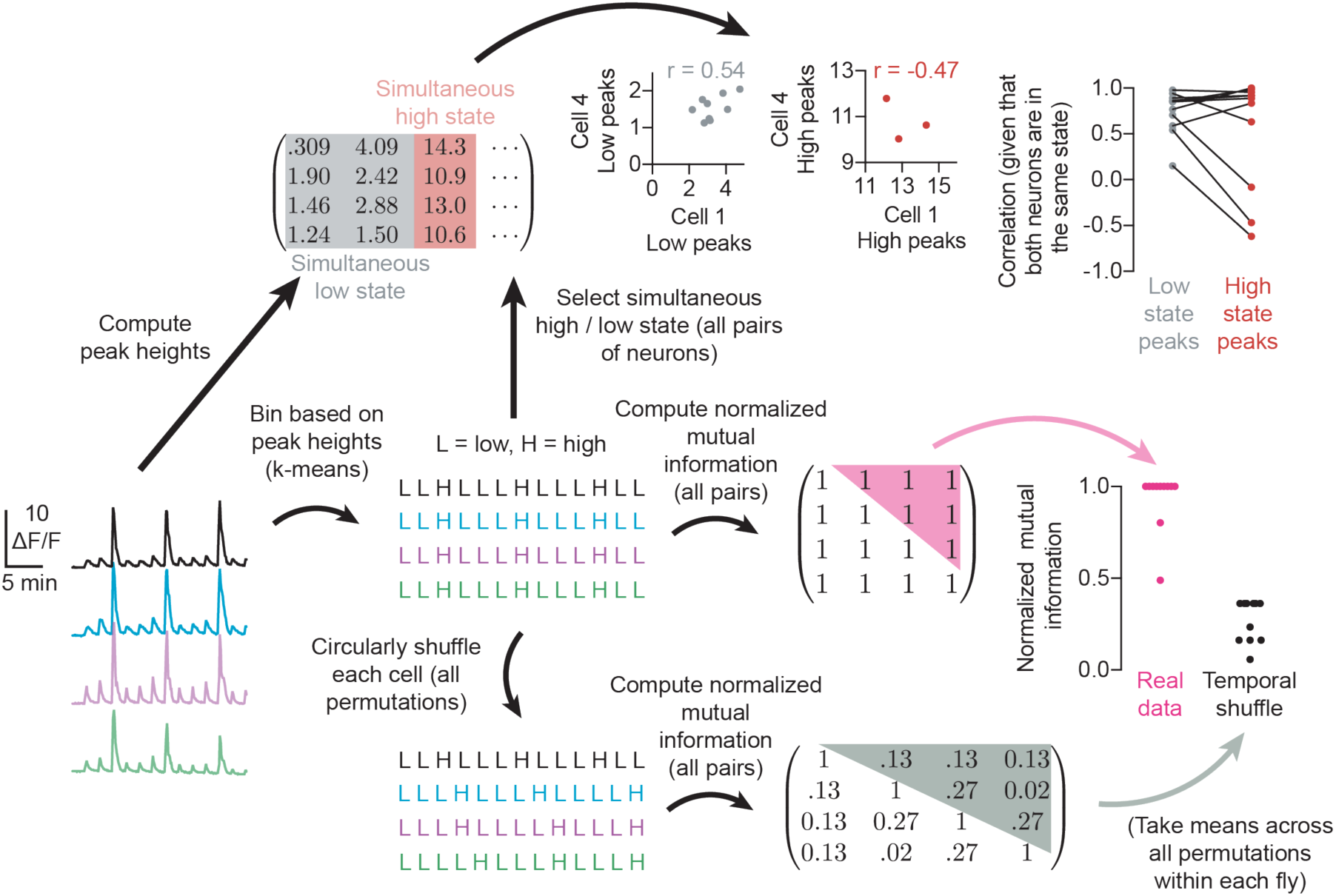
Calculating mutual information of calcium transients in response to cAMP synthesis for quantitatively identifying synchrony. The peak response in the raw fluorescence data to each optogenetic impulse was binned into two bins, either a high response or a low response, by using 2-means clustering. The mutual information of this binarized classification was computed across all stimulations, and was compared to the mutual information of a circularly permuted distribution of high and low classifications (to preserve the tendency of multiple low peaks to precede each high peak, but allow the timing of each high peak for each cell to be misaligned to that of the others). The absolute magnitude of the low peaks was highly correlated, likely due to the recurrence of the Crz neurons, while the magnitude of the size of eruptions was sometimes less correlated.

**Figure S10:**
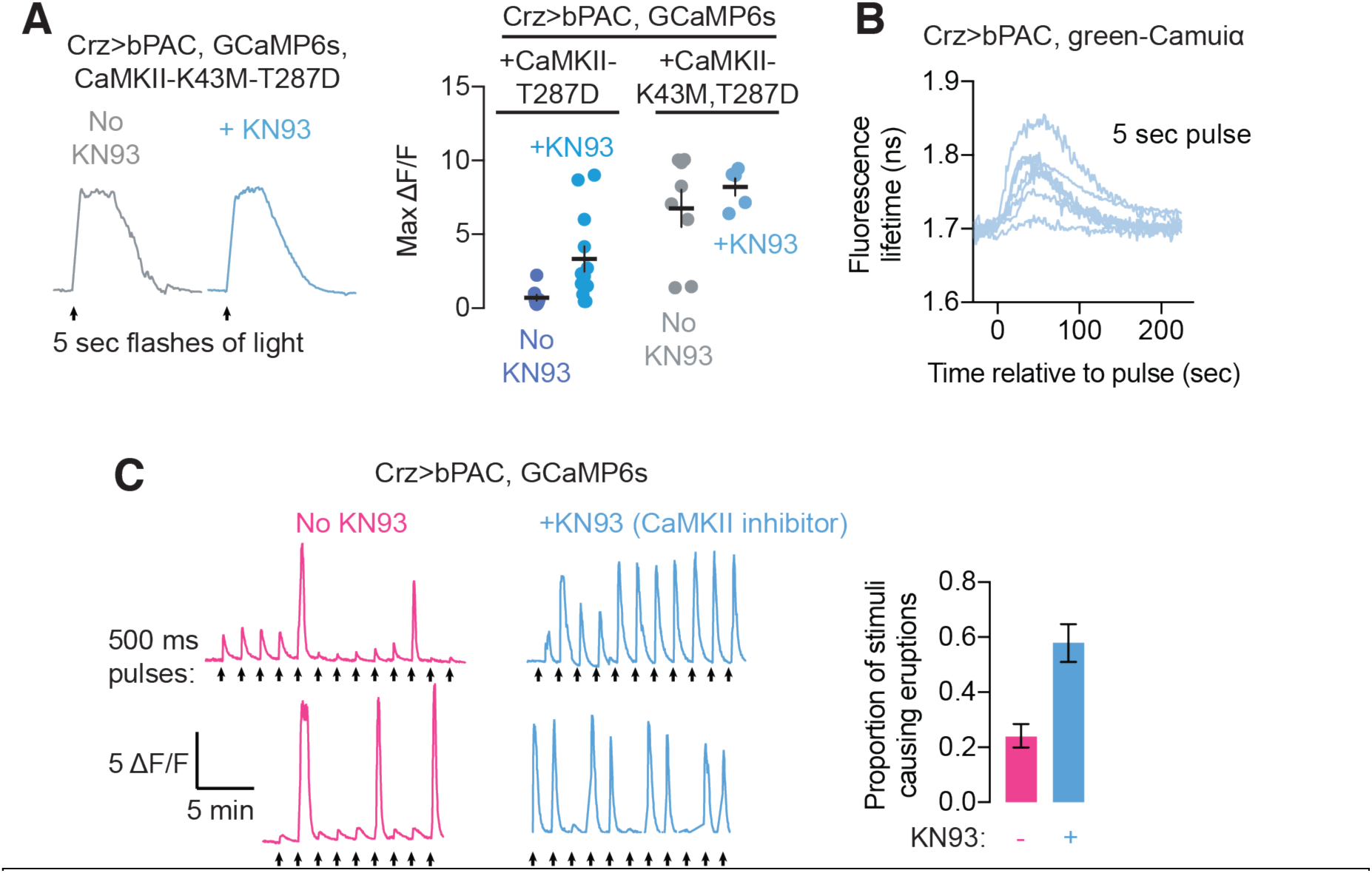
CaMKII activation after an eruption resets the Crz network. **A)** Control and summary data for Figure 6F. Application of the CaMKII inhibitor KN93 has little effect on the eruption itself in the absence of constitutively active CaMKII. **B)** Evoking an eruption with 5 seconds of stimulation of bPAC activates CaMKII as measured by green-Camuiα fluorescence lifetime. The elevation of CaMKII activity is much weaker, and so lasts a shorter duration, than with optogenetic stimulation ^6^, suggesting that different sources of intracellular calcium can be more or less effective at driving CaMKII activation, even if they result in similar overall levels of calcium elevation. **C)** Application of KN93 permits eruptions to be elicited one after another, even if pulses of bPAC activation are spaced out by 100 seconds, arguing that CaMKII activation clears out cAMP levels after each eruption.

**Figure S11:**
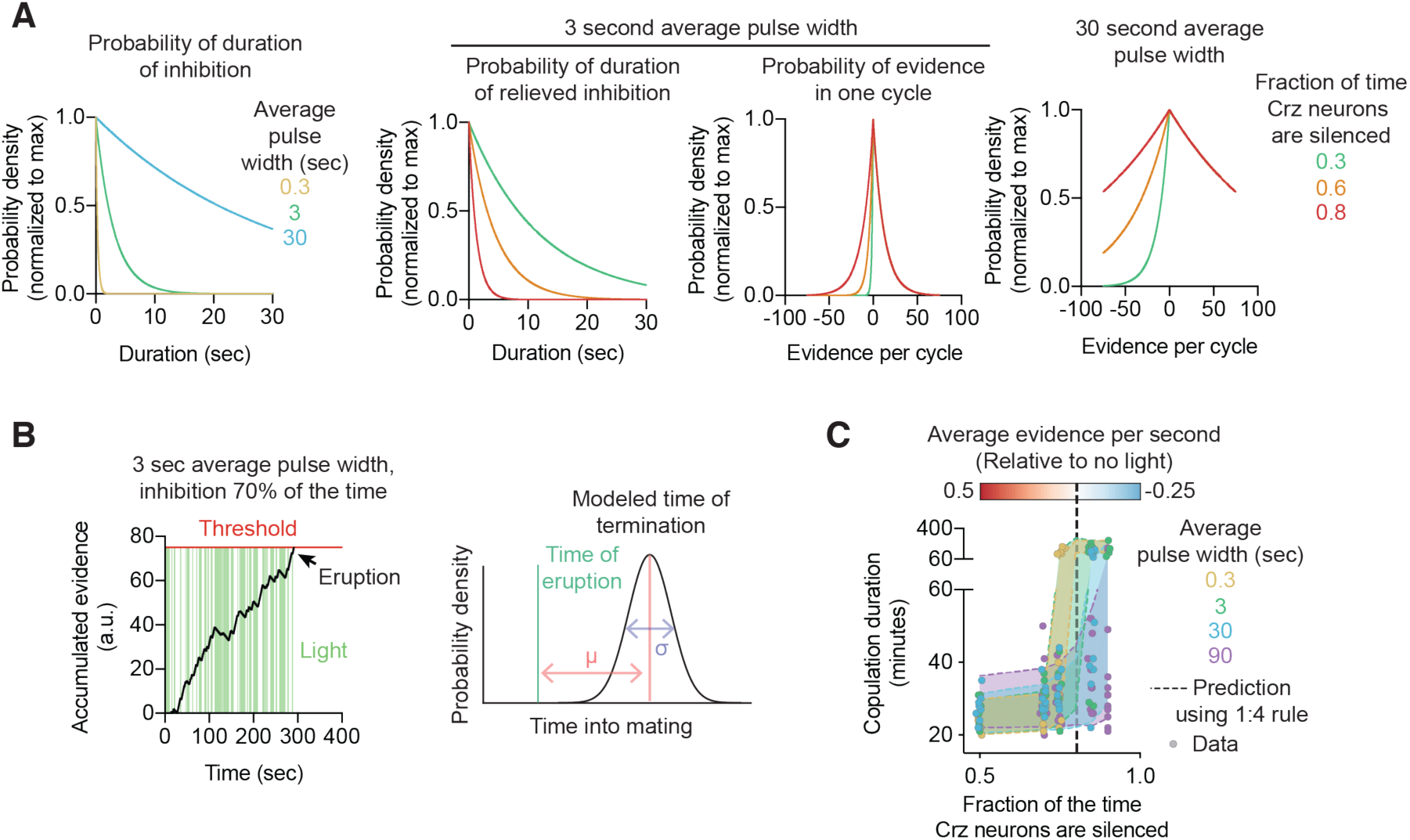
Details of randomized inhibition experiments. **A)** Schematic of the distribution of light pulses and relieved inhibition in the experiments in Figure 7E. Left: Probability of each light pulse width is determined by a single parameter: the mean width of a light pulse *m*. Pulse width is drawn from an exponential distribution with this mean: 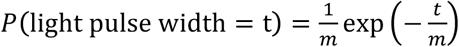 Right: Average duration of relieved inhibition is determined by the mean pulse width *m* and the fraction of the time the Crz neurons are silenced 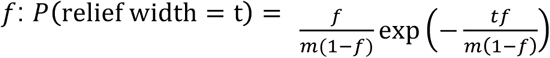 **B)** Schematized evidence accumulation to threshold to produce a simulated copulation duration. Evidence is accumulated to a threshold (equivalent to 75 seconds of no inhibition), after which a delay until mating termination is drawn from a Gaussian with mean 18 minutes. **C)** Comparison between simulated copulation durations as in **Figure S11B** and actual copulation durations. Mean pulse width is indicated by color, while position on the x-axis describes the fraction of the time the Crz neurons are silenced. Dotted lines indicate the 5^th^-95^th^ percentile copulation durations of simulated data. Individual circles show the copulation durations of single flies.

## SUPPLEMENTARY VIDEOS

**Supplementary Video 1 | The Crz neurons require over 60 seconds of electrical activity to permit a normal copulation duration**

Flies expressing GtACR1 in the Crz neurons are exposed to green light until 10 minutes into mating, when inhibition is relieved for either 30 seconds (top row) or 90 seconds (bottom row). All flies exposed to 90 seconds of relief terminate mating around 18 minutes later, while those exposed to 30 seconds of relief all persist in mating for much longer.

**Supplementary Video 2 | Calcium levels return to baseline within 100 seconds after bPAC-mediated cAMP synthesis**

Flies expressing bPAC in the Crz neurons exposed to 500 ms pulses of blue light (yellow flashes) show modest increases in intracellular calcium in response to induction of cAMP synthesis before an eruption, a massive increase in calcium levels. Left: pixelwise fluorescence, right: ΔF/F across the entire image. (Sped up ∼16x)

**Supplementary Video 3 | Eruptions are a network phenomenon and synchronous across all Crz neurons**

Flies expressing bPAC in the Crz neurons exposed to 500 ms pulses of blue light (yellow flashes). All neurons erupt synchronously. Left: Pixelwise fluorescence. Right: ΔF/F in each soma plotted in a separate color. (Sped up ∼60x).

## METHODS

### Fly stocks

Flies were maintained on conventional cornmeal-agar-molasses medium under 12-hour light/12-hour dark cycles at 25 °C. Unless otherwise stated, males were collected 0-8 days after eclosion and group-housed away from females for at least 3 days before testing. Flies expressing CsChrimson, GtACR1 and all experimental controls for experiments involving CsChrimson or GtACR1 were housed with rehydrated potato food (Carolina Bio Supply Formula 4-24 Instant Drosophila Medium, Blue) coated with all-trans-retinal (Sigma Aldrich R2500) diluted to 50 mM in ethanol for at least 3 days, unless marked as “no retinal.” These vials were kept inside aluminum foil sheaths or 3D-printed plastic cases to prevent degradation of the retinal due to light exposure. Virgin females used as partners for copulation assays were generated by heat-shocking a UAS-CsChrimson-mVenus stock (Bloomington stock #55135) with a hs-hid transgene integrated on the Y-chromosome in a 37 °C water bath for 90 minutes. This stock was selected for mating partners because the females are highly receptive to courtship, resulting in a large number of mating pairs shortly after initiation of assays, and because it has been shown that copulation duration is robust to variations in the female’s genetic background ^62^. Virgins were group-housed for 3-13 days before use. Experiments with optogenetic tools were not performed at specific times relative to the light-dark cycle of the incubator because these animals were housed in constant dark conditions for at least a day prior to the assay to preserve all-trans-retinal integrity. We did not observe any dependency of time of day on any of the behaviors described here. Detailed genotypes of all strains used in the paper are listed in a separate Table.

### Evaluation of mating

A pair of flies was scored as “mating” when they adopted a stereotyped mating posture for at least 30 seconds. This posture consists of the male mounting the female and propping himself up on her abdomen using his forelegs, while curling his own abdomen and keeping the genitalia in contact. The posture is starkly different from anything exhibited during other naturalistic behaviors and is rarely, if ever, sustained for 30 seconds during unsuccessful attempts to initiate a mating. If the flies are physically pulled apart without disengaging the genitalia (such as if the female falls or if they collide with an obstacle), the male is able to climb back into place. This posture is also maintained in the presence of threats, unless the male elects to terminate the mating, differentiating it from a “stuck” phenotype. When stuck, the male dismounts the female, orients himself away from her, and attempts to walk away, but cannot decouple their genitalia. In the rare cases when we see a male become stuck in response to a threat, we record the copulation as having terminated. Occasionally in extremely long (>1hr) mating males, the male will become stuck, possibly because the seminal fluids harden and adhere the flies together. In this case too, the onset of the stuck posture is scored as the end of mating.

### Optogenetic stimulation during behavior

#### Non-screen experiments

For CsChrimson experiments: One male and one virgin female fly were placed in each of .86” diameter 1/8” thick acrylic wells sitting 4” above 655 nm LEDs (Luxeon Rebel, Deep Red, LXM3-PD01-0350) driven using 700 mA constant current drivers (LuxDrive BuckPuck, 03021-D-E-700) and passed through frosted collimating optics (Carclo #10124). This spot of light was scattered using a thin diffuser film (Inventables, 23114-01) under the wells to ensure a uniform light intensity of ∼0.1 mW/mm^2^. The LEDs were controlled using an Arduino Mega2560 (Adafruit) running a custom script, which itself was controlled by a Raspberry Pi (either 2 or 3, running Raspbian, a Debian variant). Flies were observed by recording from above using the Raspberry Pi with a Raspberry Pi NoIR camera (Adafruit) and infrared illumination from below using IR LED arrays (Crazy Cart 48-LED CCTV Ir Infrared Night Vision Illuminator reflected off the bottom of the box) while streaming the video to a computer for observation.

For ACR experiments: The set-up was as above except using the green Luxeon Rebel, LXML-PM01-0100, and a pulse-width modulated signal to set the time-average intensity of the light to ∼5 μW/mm^2^ (approximately six times brighter than the ambient light) unless otherwise noted.

For bPAC experiments: The set-up was as above, but additionally included a blue (470 nm) Luxeon Rebel (LXML-PB01-0040) with an average intensity within the well of ∼0.25 mW/mm2. The LEDs were arranged in a “3-UP” configuration (LEDSupply) and light was passed through a frosted collimating lens (Carclo 10511)

#### During the screen

Individual pairs of males and females were placed in single wells of 32 well arenas (see Thornquist *et al.* ^6^) recorded from a height of ∼9” using a Canon camera (VIXIA HF R600). Arenas were illuminated from below using a diffuse white light source (Artograph LightPad 930 LX) with a nominal illuminance of ∼5000 lux. The light pads were controlled with an Arduino ATMEGA2560 running custom Arduino code available at https://www.github.com/CrickmoreRoguljaLabs/.

#### Heat threat experiments

Heat threat experiments were performed as in ^6^:

A similar device to the one described above was constructed, with the addition of a ¼” thick water bath underneath each well. Room temperature water was continually passed through this bath, except when heat threat manipulations occurred, when water of the temperature described in each experiment was used (controlled by a separate stopcock for each well). The LEDs above were driven with 1A BuckPucks controlled by a pulse-width modulated signal selected to ensure the average intensity of illumination is the same as in the other behavioral experiments (∼0.1 mW/mm^2^ for red light, ∼ 5 µW/mm^2^ for green, 0.25 mW/mm^2^ for blue) despite having to pass through the water

### Imaging and FLIM

Images were acquired using a modified Thorlabs Bergamo II. Samples were excited using a Coherent Chameleon Vision II Ti:Sapphire laser emitting a 920 (GCaMP6s, green-Camuiα, and FLIM-AKAR) or 1000 (cADDis_Red-Up_, cADDis_Green-Down_, and experiments performing multi-color imaging) nm beam and emission was detected using cooled Hamamatsu H7422P-40 GaAsP photomultiplier tubes, with light collected through a 16x water immersion objective (Olympus). The PMT signal was amplified using Becker-Hickl fast PMT amplifiers (HFAC-26) and passed to a PicoQuant TimeHarp 260 photon counting board, which was synchronized to the laser emission by a photodiode (Thorlabs DET110A2) inverted using a fast inverter (Becker-Hickl A-PPI-D). The TimeHarp signal was acquired by custom software (FLIMage, Florida Lifetime Imaging) which was also used to control the microscope. For intensity imaging, all detected photons within a pixel were summed together, regardless of arrival time relative to the excitation pulse. Optogenetic stimulation was performed by excitation with a blue LED (Thorlabs M470L4) through a liquid light guide (Thorlabs LLG5-8H) for an incident intensity at the sample of ∼XXX. 128×128 pixel images were acquired at a rate of ∼4 Hz.

#### Calcium and cAMP imaging quantification

Most images were processed by simply taking the sum of all photon counts across the image, and then subtracting the background (estimated as the median pixel value, which was almost always 0 photon counts). Because our driver does not label any other neurons in the abdominal ganglion, we did not need to use ROIs or image segmentation to restrict our analysis to specific pixels. **Δ*F/F*** was computed as 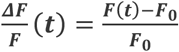 where 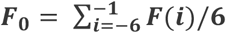 corresponding to the first frame of optogenetic excitation. Analysis was performed using custom Python3 code.

When analyzing multiple neurons simultaneously, rectangular ROIs were drawn around identified cells or processes and the same analysis as above was performed.

Experiments using only GCaMP6s used a 920 nm excitation wavelength, while those performed in conjunction with cADDis_Red-Up_ used a 1000 nm excitation beam. Experiments using cADDis_Green-Down_ also used a 1000 nm excitation wavelength.

#### FLIM quantification

FLIM data was collected for each pixel in 64 time bins of width 200 picoseconds. All pixels across the series of images were pooled to fit the parameters of a multiple-time constant model Exp(t) convolved with the instrument response function IRF(t) (assumed to be a Gaussian) by finding the maximum likelihood estimate of the parameter values. The probability that a photon will be detected at time t is then described by the relation:

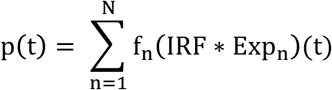

where f_n_ refers to the proportion of excited fluorophores in state n with time constant τ_n_. The two convolved distributions are

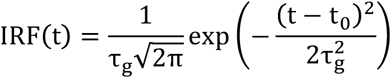

corresponding to the probability of a given instrumentation induced shift in the detected arrival time and

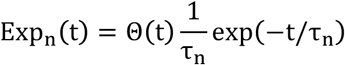

with Θ(t) the Heaviside theta function Θ(t) = 1 if t ≥ 0 and 0 otherwise. This corresponds to the probability of an excited fluorophore in state n emitting a photon at time t. The convolution of these two distributions is

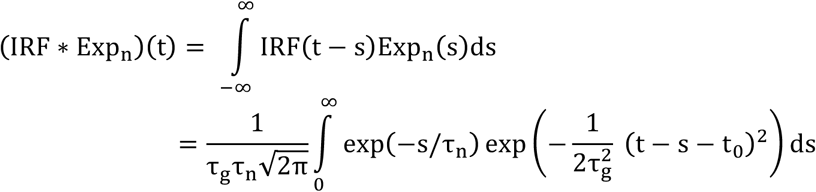

We can evaluate this integral by completing the square inside the exponential:

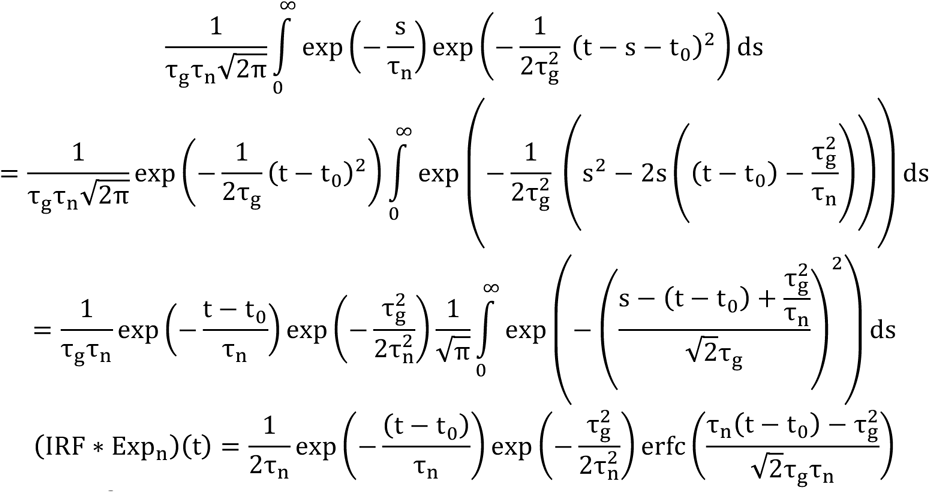

with 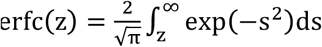 the complementary error function. We then fit the parameters {τ_n_}, {f_n_}, {τ_g_}, {t_0_} by minimizing the log-likelihood function ℒℒ ({n_i_}) (with n_i_ the number of counts in photon detector time bin i)

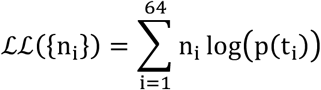

with t_i_ corresponding to the time of detector time bin i, subject to the constraints ∑_n_f_n_ = 1, 0 ≤ f_n_ ≤ 1, 0 < τ_g_, t_0_, τ_n_.

We used the obtained estimate for t_0_ to compute the “empirical lifetime” τ, the mean arrival time of a photon ^12^. τ is defined as

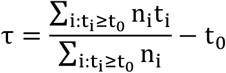

The typical value for t_0_ was approximately 2.9+/-0.1 nanoseconds. τ_g_ values ranged from 0.1 to 0.2 nanoseconds. Other parameters were used to be sure fluorophores weren’t forming aggregates, in which their fit parameters would deviate from those typically observed. Usual values for τ_1_ and τ_2_ for various fluorophores were as follows: green-Camuiα: 0.6, 2.2 nanoseconds; FLIM-AKAR: 0.7, 2.1 nanoseconds; cADDis_Green-Down_: 1.2, 2.8 nanoseconds, GCaMP6s: 2.8 nanoseconds (usually monoexponential).

#### Dissections

Flies used in imaging experiments were dissected in chilled saline (103 mM NaCl, 3 mM KCl, 5 mM TES, 8 mM trehalose, 10 mM glucose, 26 mM NaHCO_3_, 1 mM NaH_2_PO_4_, 3 mM MgCl_2_, 1.5 mM CaCl_2_, pH ∼7.25, 270-275 mOsm), as in our previous work ^6^. The same saline solution was used during imaging experiments.

### Antibodies and immunohistochemistry

All samples were fixed in PBS with Triton X-100 and 4% paraformaldehyde for 20 minutes, then washed three times with PBS/Triton X-100 for 20 minutes each before application of antibodies. All samples were incubated with the primary antibody for two days, washed three times with PBS/ITriton X-100 for 20 minutes each, incubated with the secondary antibody for two days, then washed three times as before and mounted on coverslips using VectaShield (Vector Labs). Antibodies used are as follows:

Chicken anti-GFP (GFP-1010, Aves Labs, 1:1000 dilution)

Rabbit anti-DsRed (323496, Clontech 1:1000 dilution)

Mouse anti-nc82 (Developmental Studies Hybridoma Bank, 1:8 dilution)

Donkey anti-chicken 488 (703-545-155, Jackson ImmunoResearch, 1:400)

Donkey anti-rabbit 555 (A-31572, Invitrogen, 1:400 dilution)

Donkey anti-mouse Cy3 (715-166-150, Jackson ImmunoResearch, 1:400 dilution)

Donkey anti-rabbit 647 (711-605-152, Jackson ImmunoResearch, 1:400 dilution)

Donkey anti-mouse 647 (711-605-151, Jackson ImmunoResearch, 1:400 dilution)

### Confocal microscopy

Confocal images were collected using a Zeiss LSM 710 through a 20x air objective (Olympus PLAN-APOCHROMAT) controlled by Zen software, and analyzed using ImageJ.

### Generation of transgenic flies

UAS-FLIM-AKAR flies were generated by amplifying the gene out of the AAV-FLIM-AKAR construct Addgene #63058 using the primers GCGGCCGCGGCTCGAGatggtgagcaagggcgaggagct and ACAAAGATCCTCTAGAttactcgatgttgtgcctgattttgaagttgac. This was then cloned into a linearized 20xUAS-IVS-mCD8::GFP (Addgene #26220) (digested with XhoI and XbaI) using InFusion and injected into attP2 and attP40 by BestGene, Inc.

### Statistics

#### General framework

Throughout this manuscript, we take a Bayesian approach to parameter estimation because it more closely corresponds to the inference procedures we are performing, and as such all reported windows and intervals correspond to the mass of the posterior distribution for the inferred model parameter. This is because the Bayesian approach corresponds to inference about the values of descriptors of our model (e.g. in our data, the probability of terminating the mating in response to some stimulus), rather than consistency of a data set with a particular value that the model might take. With this approach, we can make statistical claims about our belief in the magnitude of effects, rather than simply reporting their deviation from that produced by a null hypothesis. We do, however, recognize that the frequentist approach is more commonplace, and so present our data in a manner that is as consistent as possible with typical frequentist reporting and hypothesis testing. We use noninformative priors ^63^, so this trivially corresponds to the usual Central Limit Theorem statistics in the case of estimating the variability of means, but a slightly different estimator on proportions. Our results and their interpretation do not hinge, in any case, on precise statistical methodology, as our effects tend to be very large, and so this decision is more philosophical than effectual.

#### Credible intervals for proportions

All proportions are modeled as Bernoulli random processes with probability *p*and presented as the sample estimate 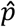 for the proportion *p* (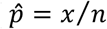 with *x* the number of observed successes and *n* the total number of observations, the maximum likelihood estimate, rather than the maximum posterior estimate, for consistency with standard data presentation). This point is surrounded by a 68% credible interval (selected to be similar to the broadly familiar SEM metric, which is itself a 68% credible interval on the mean under a uniform prior), generated by sampling from the posterior distribution using Markov Chain Monte Carlo (MCMC, Metropolis-Hastings algorithm) with the noninformative Jeffreys prior 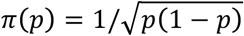 and selecting the 16-84% window of this empirical estimate of the posterior. The Jeffreys prior was selected because it gives a posterior that is in a sense invariant under reparameterizations ^64^, and thus gives a consistent result between our posterior distributions even when we invert or transform the inference problem. The window generated by this method is also a numerical approximation of a 68% confidence interval with the corresponding frequentist properties.

#### Determination of long mating or normal mating

As in Thornquist *et al.*^6^, we estimated the probability of a given experimental condition producing a long mating duration using a hierarchical Bayesian model:

For a given time *t*, we estimated the posterior distribution of the parameter *p*of a Bernoulli process, which was used to select the posterior distributions used to sample the mean and variance of the Gaussian from which an individual fly’s copulation duration was drawn. Explicitly:

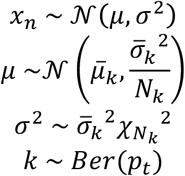

with 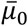 and 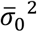 the mean and variance of the “no light” condition, and 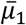 and 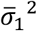 the mean and variance of the “light at start” condition, and likewise *N_k_* the corresponding size of those samples. The prior on *p_t_* was a Beta distribution with *α* = *β* = 0.5, the Jeffrey’s prior, and we used MCMC to estimate the posterior distribution of *p_t_*. We then plotted the maximum *a posteriori* value for *p_t_* along with 68% credible intervals.

A similar model was used for the “relief from inhibition” experiments in which inhibition was relieved after 10 minutes of mating. These flies would mate for ∼18 minutes longer (in contrast to those flies in which inhibition was not relieved, that would typically mate for at least another hour). The flies in which inhibition was removed at 10 minutes were used instead of the “no light” condition for generating the priors on 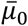 and 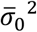 as in Figure 7A.

#### Computing normalized mutual information of eruption timing

For each cell, we found the peak response to each pulse of light and used these to populate an *m* × *n* dimensional matrix *F*, with *m* cells for each of the *n* pulses. Each row of the matrix was fit to a Gaussian mixture model with two components, and then each pulse response was classified as having most likely been produced by one of the two Gaussians. Those in the cluster with the highest mean were labeled “eruptions” and those in the other cluster were labeled “non-eruptions.” We then measured the mutual information using the *n* columns ***x*** ∈ {0,1}*^m^* to estimate the mutual information *M*(*x_i_*; *x_j_*) with *x_i_* the *i*^th^ row of each ***x***. The mutual information was normalized to create the matrix

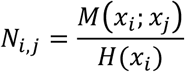

and the upper triangle (excluding the diagonal) of this matrix (i.e. the normalized mutual information between each pair of cells) was plotted in **Figure 5C**. This was compared to all permutations of the above in which each row of *F* was circularly permutated (that is, *F* was replaced by *F*^’^ where 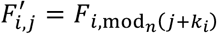 with a different *k_i_* for each row) to generate a shuffled sequence that had the same overall patterning as *F* but in which the relationship between peak responses across cells is scrambled.

All code for these analyses (and corresponding hypothesis testing) was written using the freely available Python packages `numpy` and `pymc` and is available on Github at https://github.com/CrickmoreRoguljaLabs.

#### Hypothesis testing on proportions

To test the hypothesis that two sample proportions were drawn from the same Bernoulli process, we used Fisher’s exact test.

#### Credible intervals on means

We use the standard SEM estimator for variability of sample means 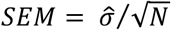 with 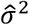 the unbiased estimator of sample variance and *N* the sample size, which corresponds to the 68% credible interval for the sample mean using the “improper” uniform prior.

#### Hypothesis testing on distributions

We use the nonparametric Mann-Whitney U test on rank sums for differences in distributions of copulation duration. We then correct for multiple comparisons by using the Holm-Bonferroni correction on our criteria for statistical significance (with the number of hypotheses being the number of unique pairs of comparisons, *n* (*n* − 1)/2).

### Modeling

#### Analysis of mutual information between elapsed time and CaMKII activity

We use the standard definition for the entropy of a random variable *s* with domain *S* and measure *ds*:

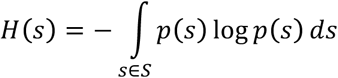

with log(*x*) referring to the logarithm base 2 (giving units of bits). We consider a variable *I*(*t*) that takes the value *I*(*t*) = 0 if *t* ≤ τ and *I*(*t*) = 1 if *t* ≤ τ with *t* ∼ *Uni*(0,10) uniformly distributed from 0 to 10. Then the entropy 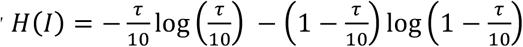

We can then define the conditional entropy

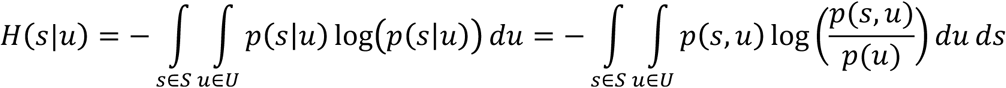

for a random variable *u* defined on domain *U* with measure *du* to compute the mutual information between *s* and *u*, i.e. the extent to which knowledge of *u* reduces the entropy of *s*:

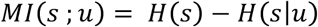

Then if the random variable *s* is determined by *t* as well, we can compute the mutual information *MI*(*I*; *s*) from *H* (*I* |*s*):

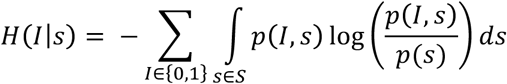

We then note that 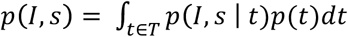 to rewrite this as:

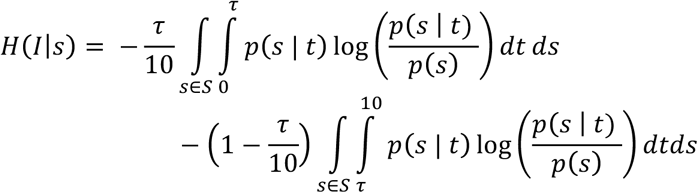

or, evaluating the integral over *t*:

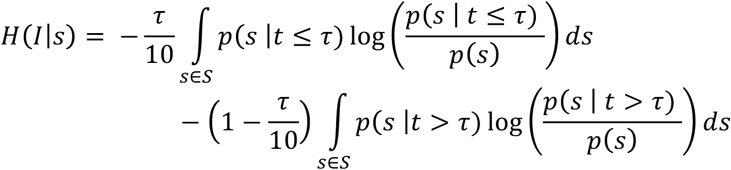

since *p*(*I* = 0 | *t* > τ) = *p*(*I* = 1 | *t* ≤ τ) = 0. For green-Camuiα, we approximated the integral by creating bins {*s_i_* } of size 0.02 nanoseconds, computed *p*(*s_i_* |*t* ≤ τ) and *p*(*s_i_* |*t* > τ) and *p*(*s_i_*) by counting the number of image frames in which the value for green-Camuiα was within *s_i_* and dividing that by the total frames being considered (e.g. all the frames such that *t* ≤ τ when computing *p*(*s_i_*|*t* ≤ τ)).

#### Analysis of mutual information between elapsed time and behavior

We use the same expression as for the analysis of green-Camuiα data, except now *s* ∈ {Long mating, normal mating}. The value of *p*(*s* |*t* > τ) was computed using the mean from the Bayesian analysis described above when τ∈{0,1,5,6,7,8,10} (in minutes), the values for which we performed experiments and a linear interpolation between those values was used for τ otherwise.

#### Evidence accumulation

We model the accumulation within cycle *t* as

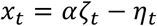

where *ζ_t_* is the positive evidence accumulated (when the light is off), *η_t_* is the negative evidence accumulated (when the light is on), and *α* is a scale factor. Both *ζ_t_* and *η_t_* are exponentially distributed, with the mean of *ζ_t_* being *μ* and the mean of *η_t_* being *μf*/(1 − *f*), with *f* the duty cycle (the proportion of a cycle, on average, that the light is on). Exponential distributions were chosen because they are the maximum entropy distribution when the mean is specified, and so in a sense are maximally disordered. More explicitly:

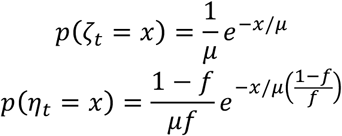

Then the evidence accumulated in a trial, *x_t_*, has the distribution

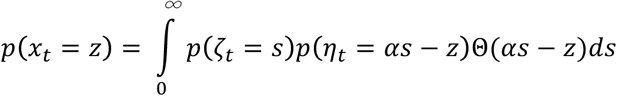

where Θ(*x*) is the Heaviside theta function, Θ *x* = 0 if *x* < 0 and Θ(*x*) = 1 otherwise.

The above can be written as

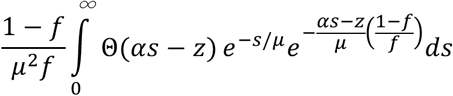

or

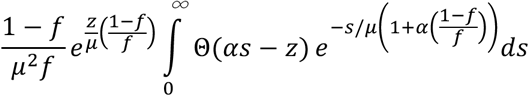

If *z* < 0, then the *Θ* is just 1 and we have that

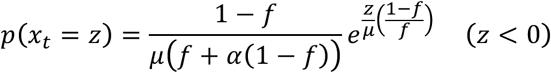

but if *z* ≥ 0 we change the bounds of the integral to find

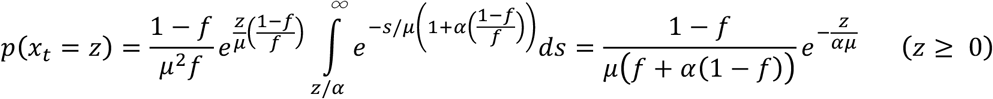

so that

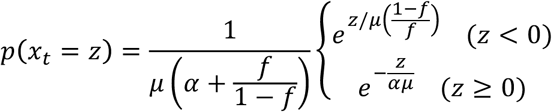

We define 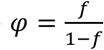 so that the above expression simplifies to

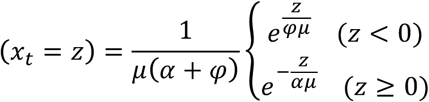

Evidence accumulation follows the model

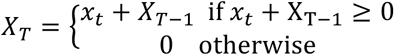

with *X_T_* the accumulated evidence at time *T* and *x_t_* i.i.d. random variables distributed as above. We then set a threshold *θ* and numerically approximated the distribution of the random variable *t* _accum_ = inf {*t* | *X_t_* ≥ *θ* } by generating 1,000 values of *x_t_* for each simulated fly according to the probability distribution above and measuring *t* _accum_ for each fly. The simulated copulation duration was estimated with:

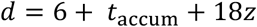

where *z* ∼ 𝒩0,9 a normal random variable estimated from the data in **Figure 7A**, 18 corresponds to the mean latency between the eruption and the termination of mating, 9 = 3^2^ the mean *σ* from **Figure 7A**, and 6 corresponds to the time for CaMKII to decay to baseline. This approximation was performed for 1000 flies for each parameter pairing *μ*,*φ* with *α* = 4 to generate the predicted distributions in **Figure 7E**. If *X_t_* < *θ* for all *t, d* was drawn from the Gaussian in **Figure 1A**, with mean 176 minutes and standard deviation 40 minutes.

### TABLES

**Supplemental Table 1.**
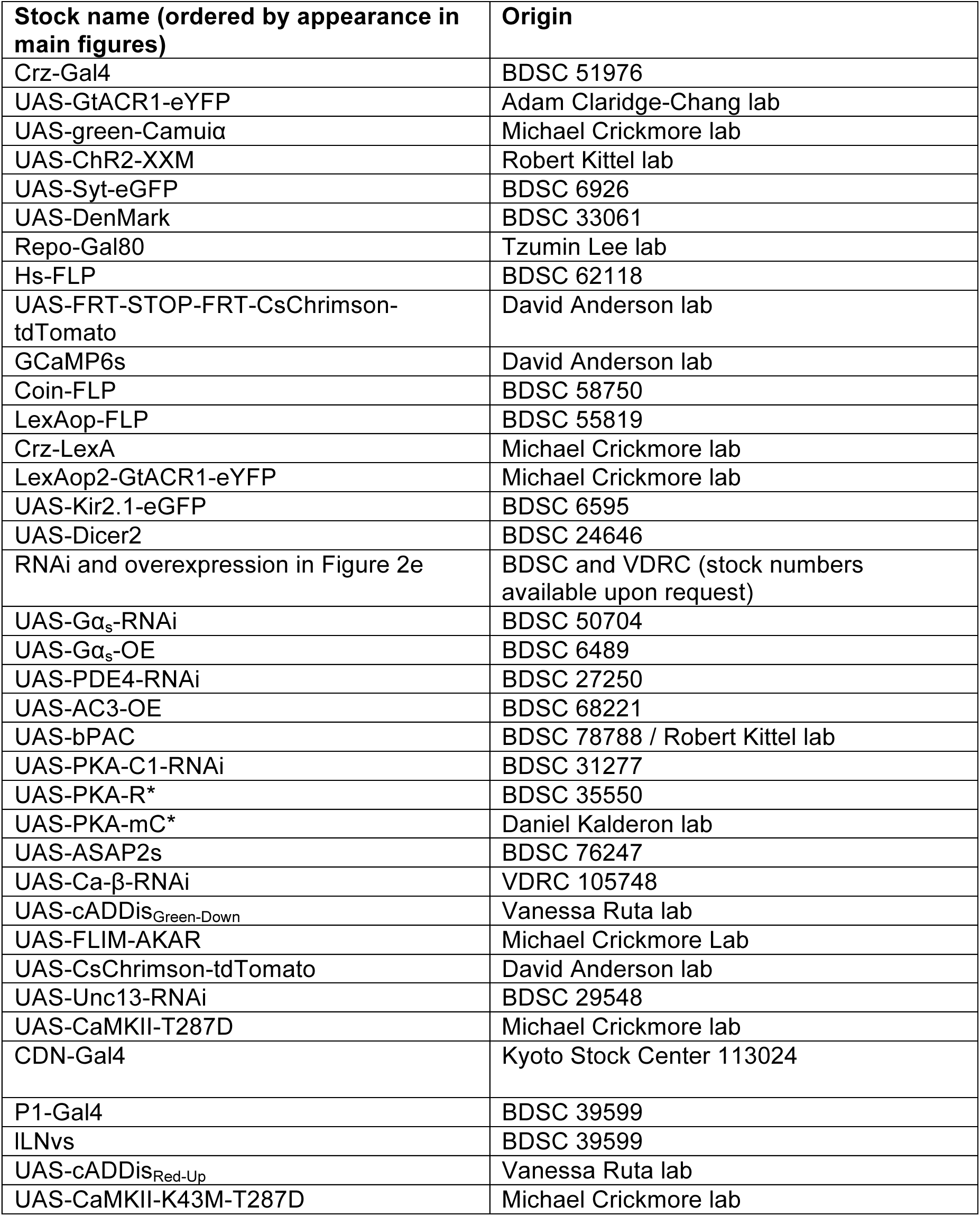
Fly stocks used and origin.

**Supplemental Table 2.**
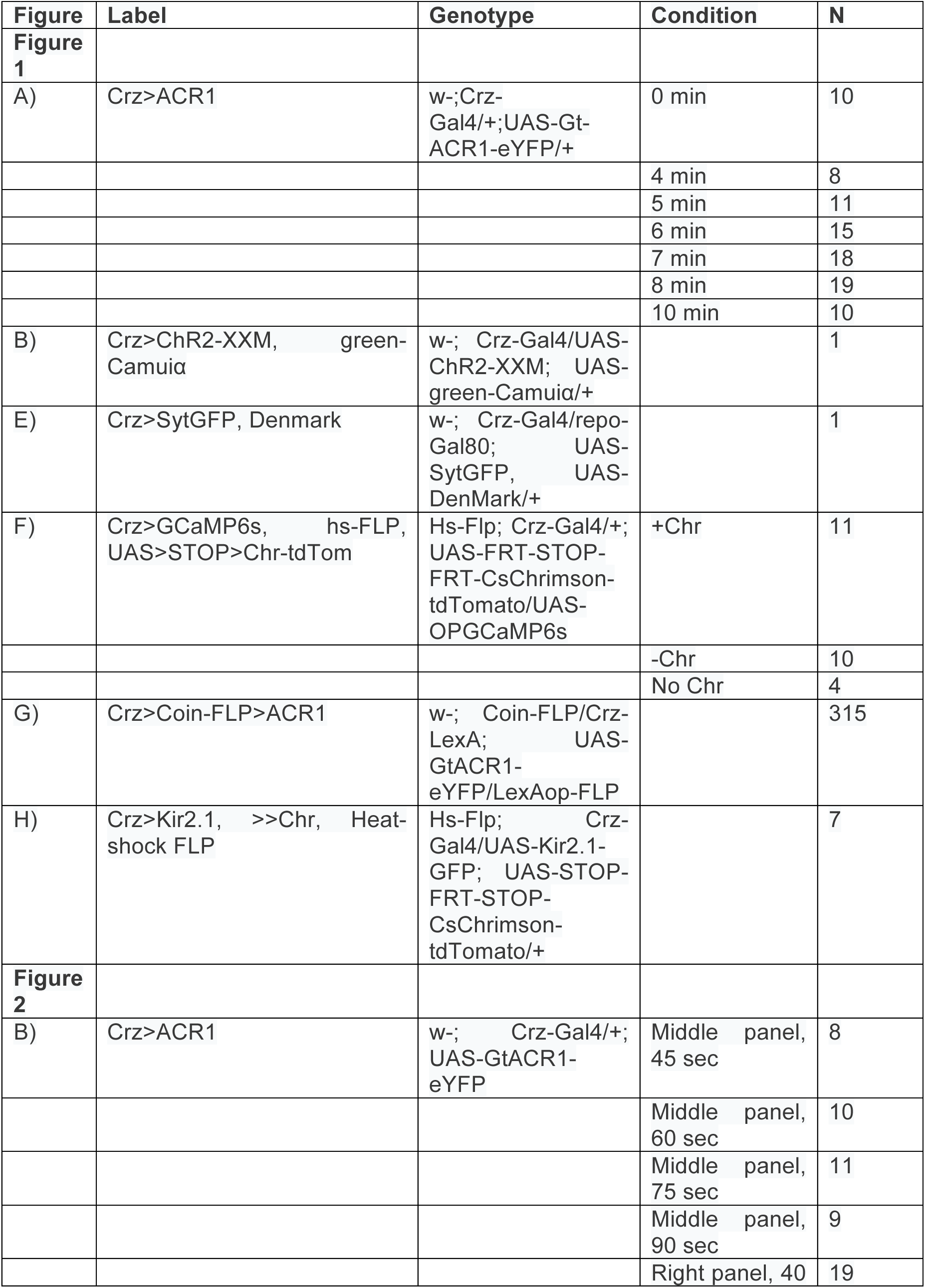
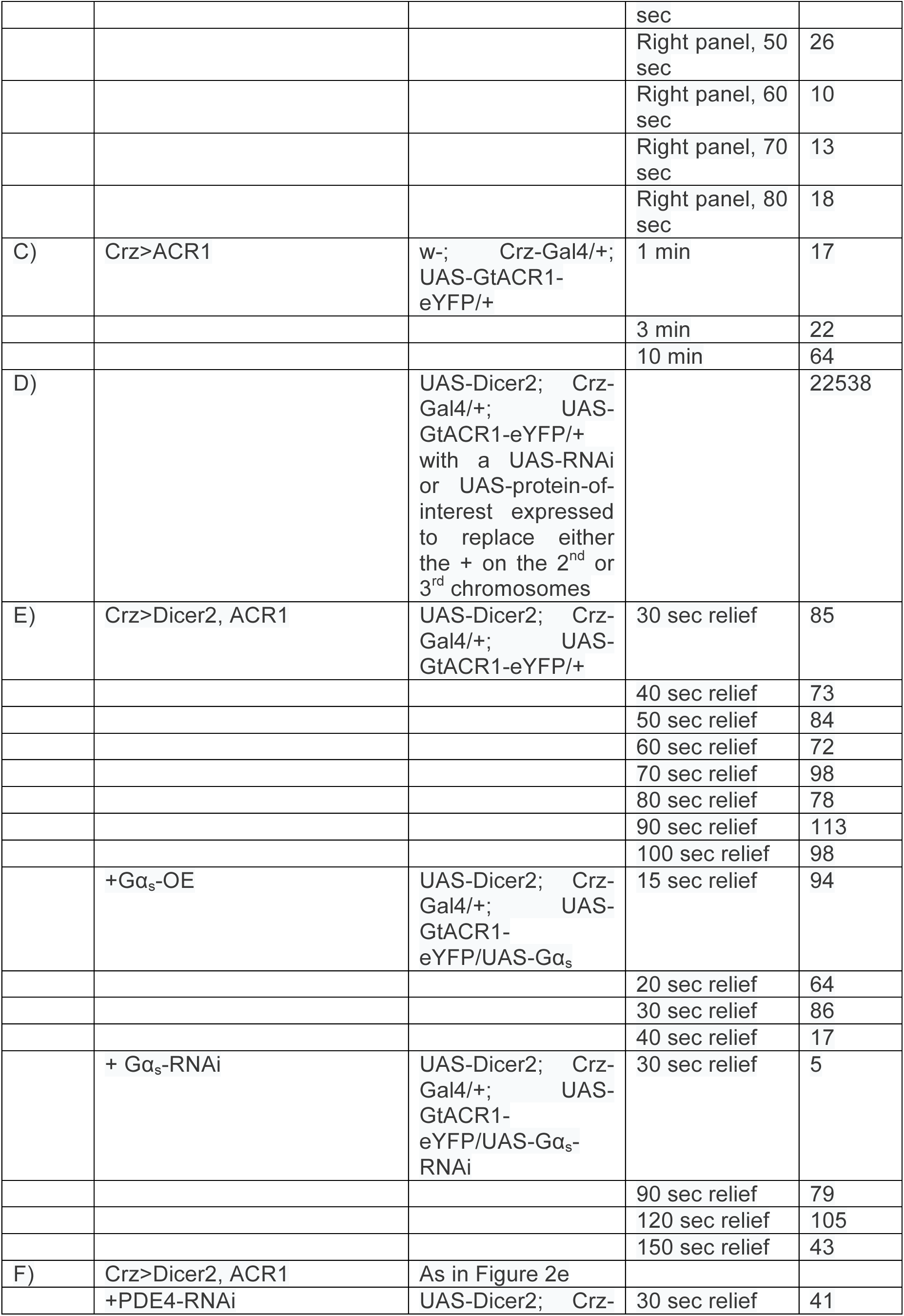
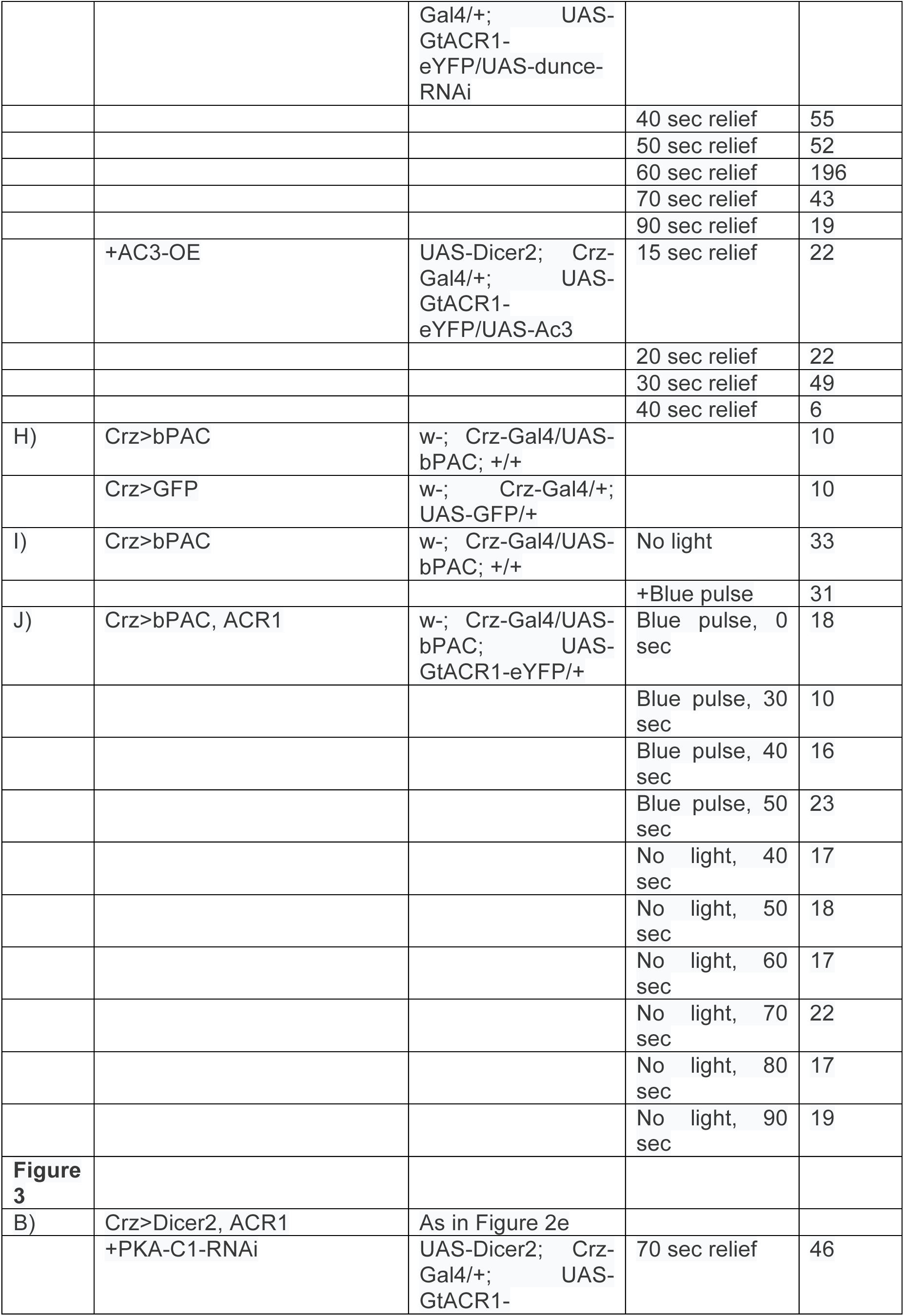
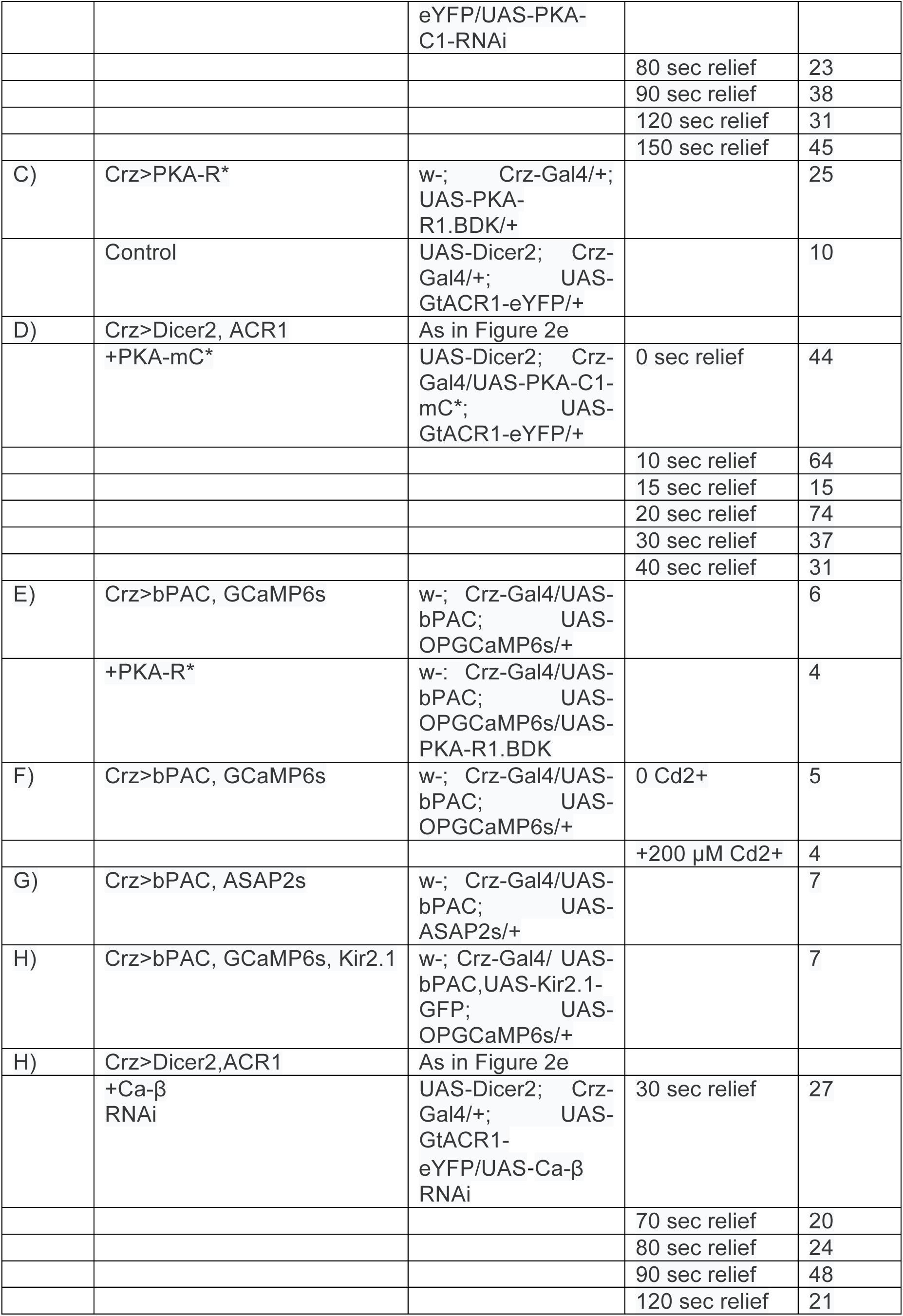
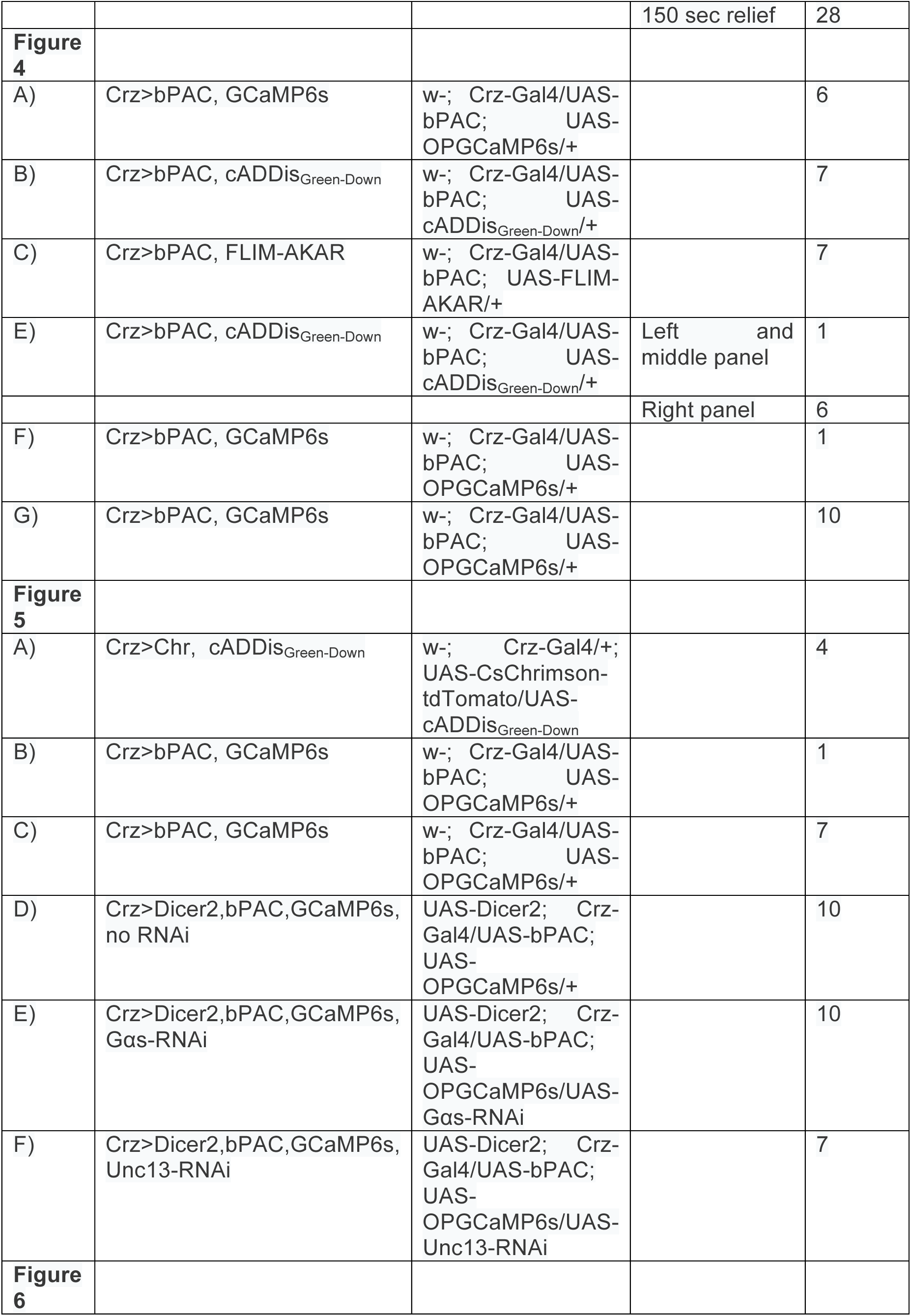
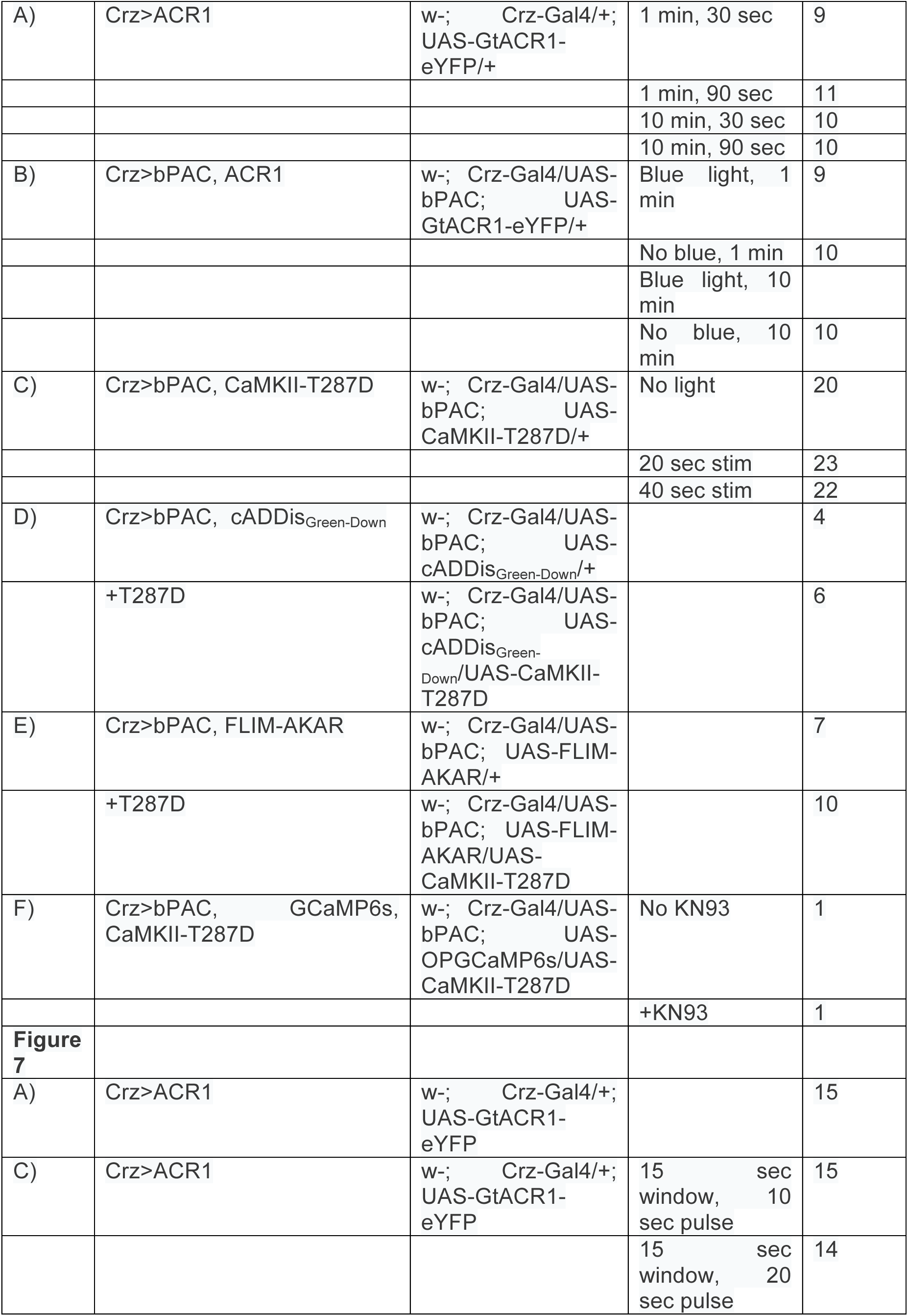
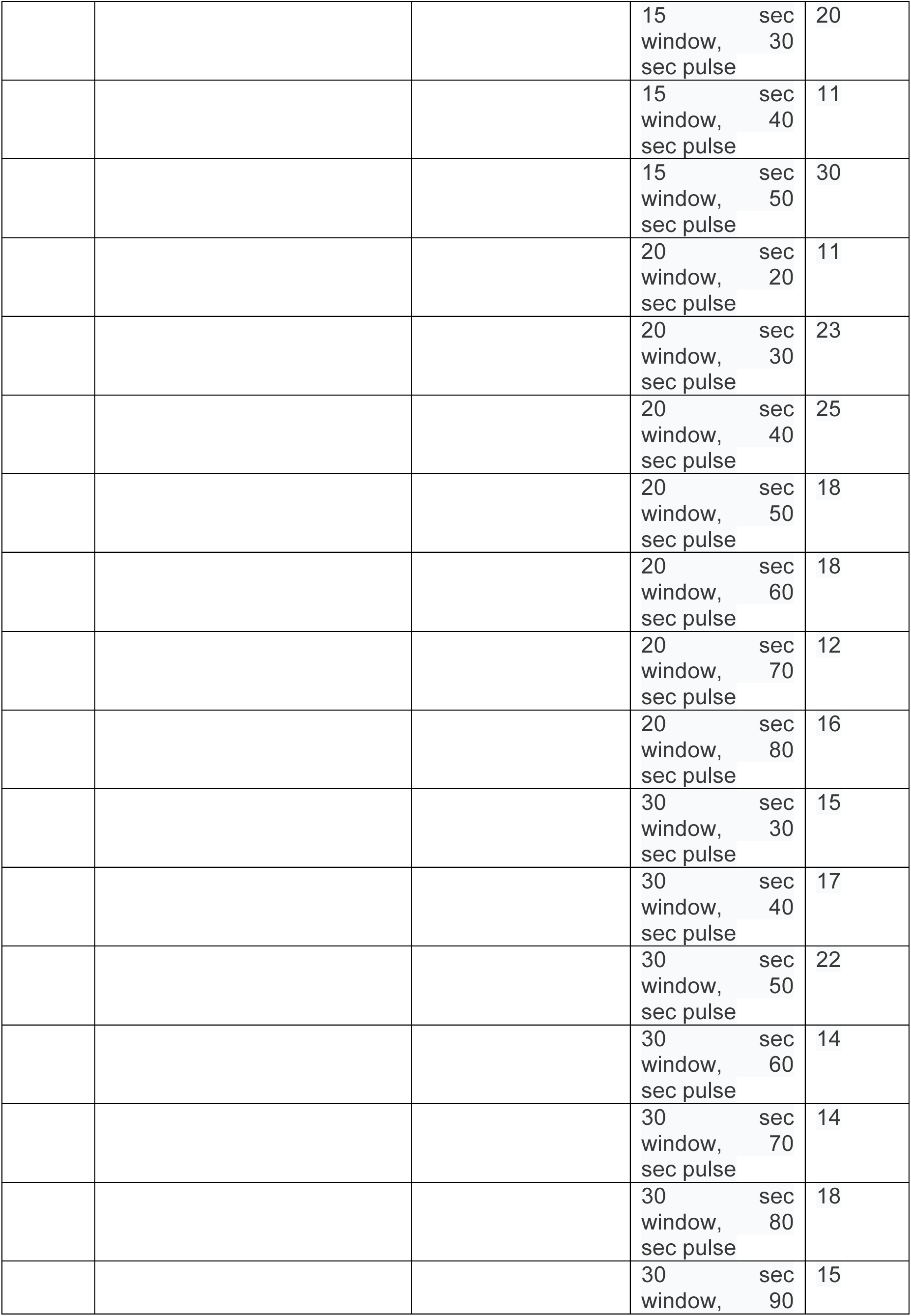
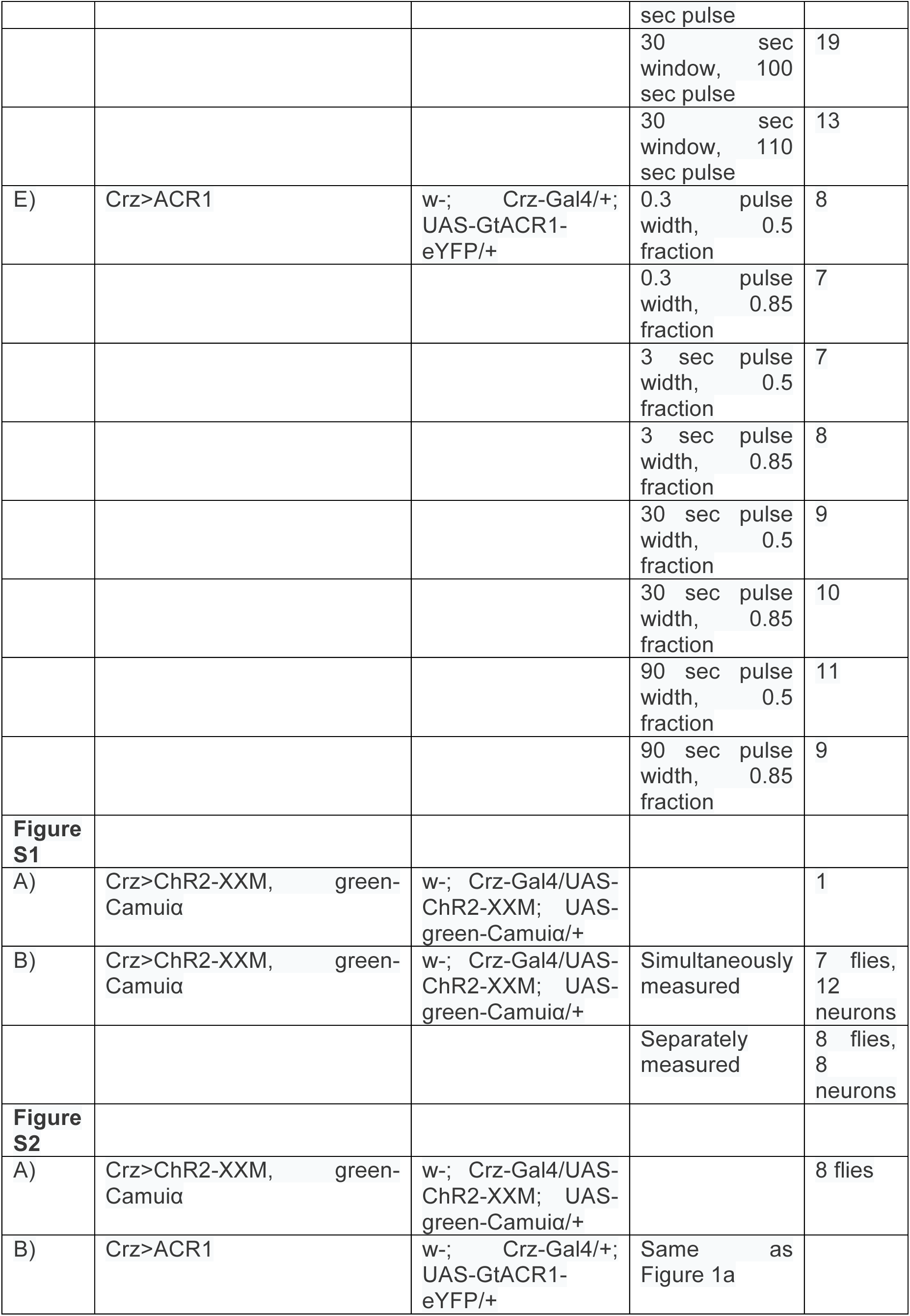
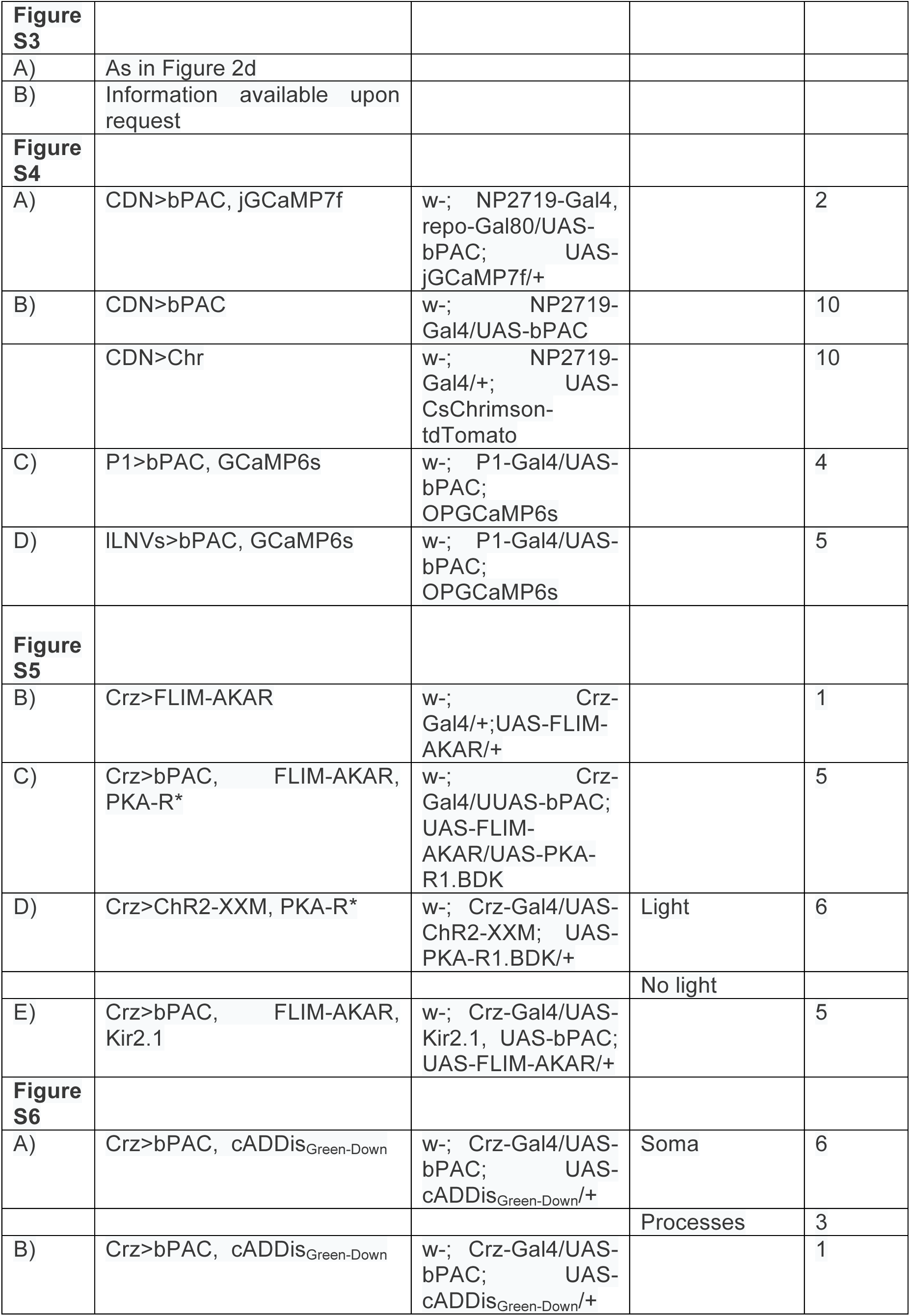
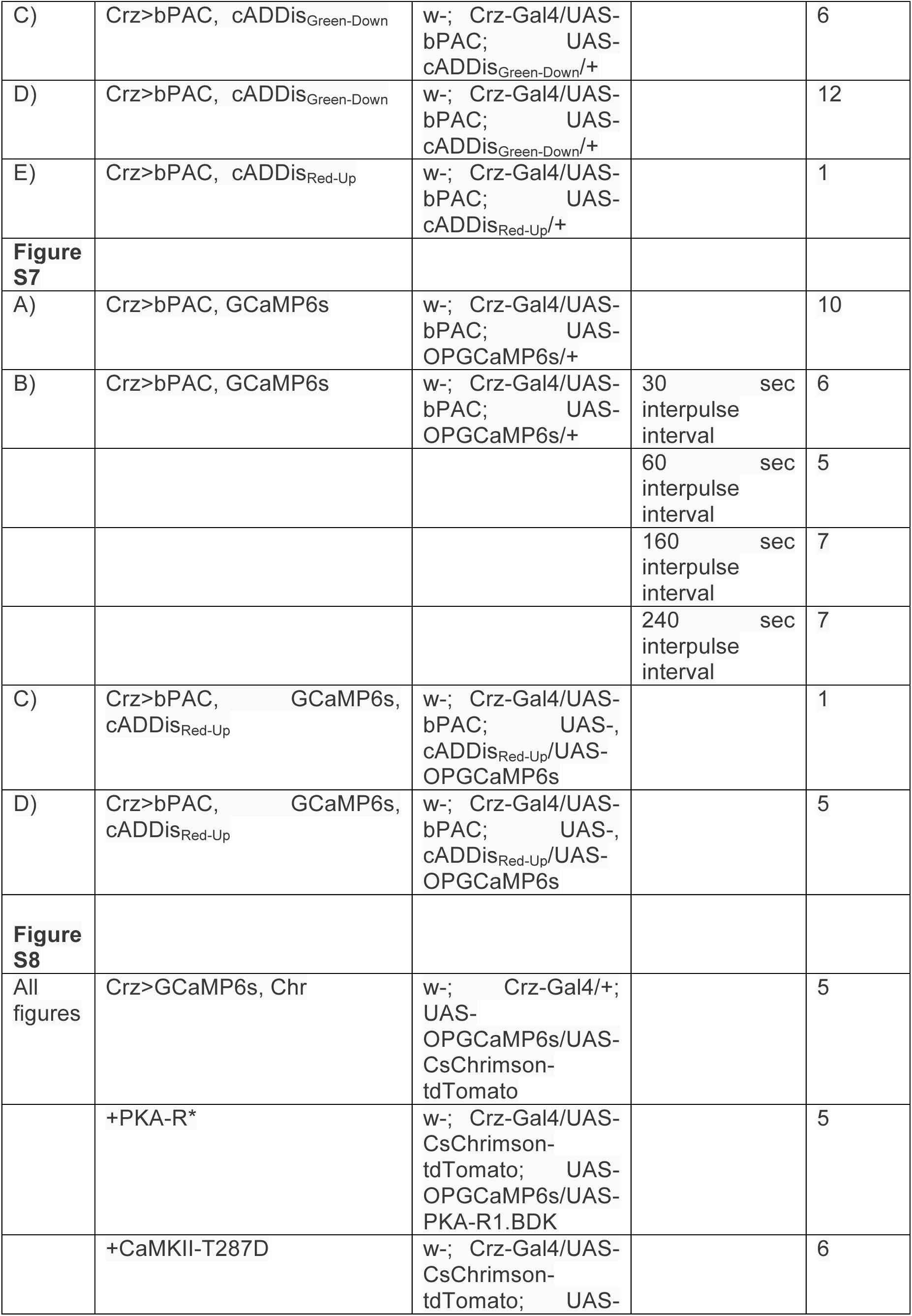
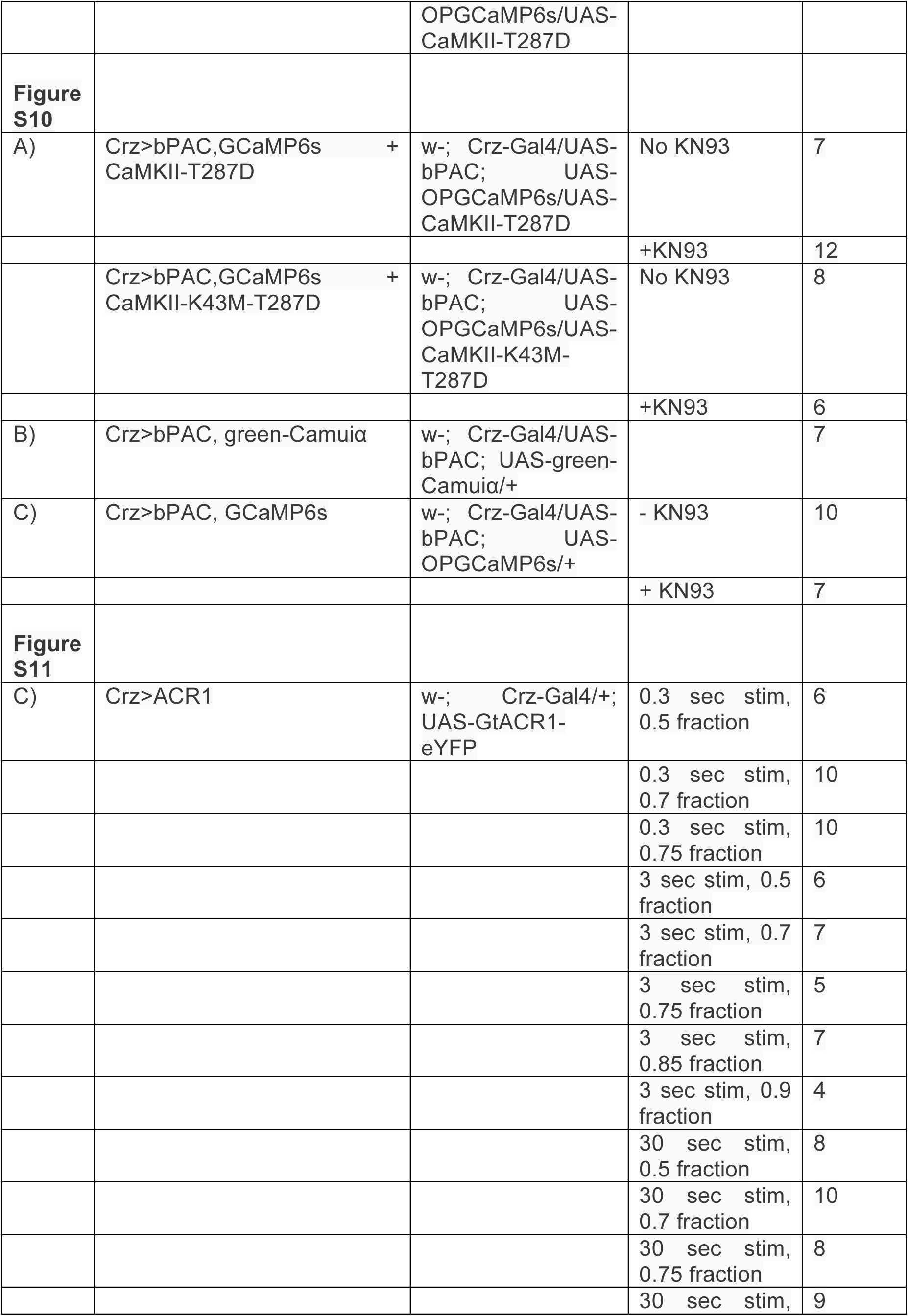
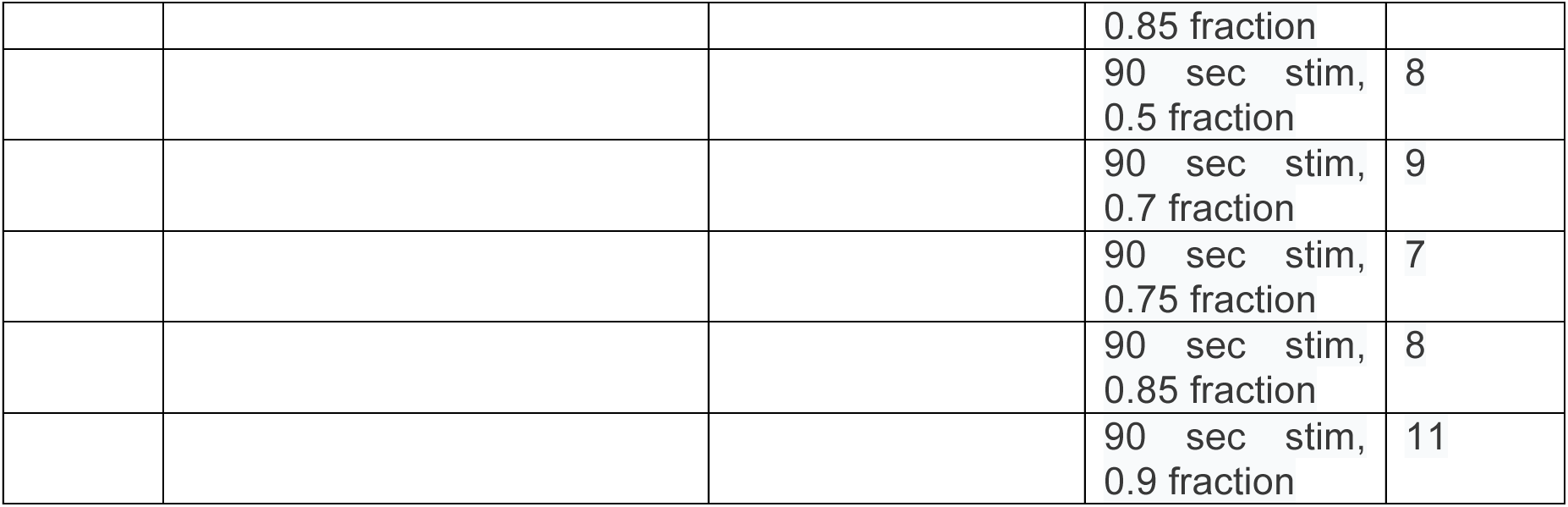
Genotypes and number of flies per experiment.

**Supplemental Table 3.** Statistical analyses and p-values.

| **Figure + Test used** | **Null hypothesis** | **p-value (significant values in red at alpha = 0.05 using Holm-Bonferroni correction for number of comparisons)**<br>**Statistical significance indicated in red** |
| --- | --- | --- |
| **Figure 1** |  |  |
| **F)** Mann-Whitney U test (groups numbered from left to right: +Chr = 1, -Chr = 2, No Chr = 3) | No difference in distributions of peak responses | 1-2: 0.0097, 1-3: 0.0015, 2-3: .0028 |
| **G)** Chi-squared test for deviation between the predicted model (silencing 3 or 4 neurons produces long matings) and the data | The data is explained by long matings resulting only from silencing 3 or 4 neurons | 0.8506 |
| **Figure 2** |  |  |
| **B)** Mann-Whitney U test (grouped numbered from left to right) | No difference in copulation durations between treatments | 1-2: 0.5681, 1-3: 0.0457, 1-4: 0.001, 2-3: >0.9999, 2-4: 0.368, 3-4: >0.999 |
| **C)** Fisher's exact test (groups numbered from left to right) | No difference in proportion long matings between treatments | 1-2: 0.0056, 1-3: <0.0001, 2-3: <0.001 |
| **H)** Fisher's exact test | No difference in proportion ejaculation | <0.0001 |
| **I)** Fisher's exact test | No difference in the proportion terminating in response to heat threats | 0.0007 |
| **Figure 3** |  |  |
| **C)** Mann-Whitney U test | No difference in copulation durations between genotypes | <0.0001 |
| **E)** Mann-Whitney U test | No difference in peak fluorescence responses | 0.0095 |
| **F)** Mann-Whitney U test | No difference in peak fluorescence responses | .0159 |
| **Figure 4** |  |  |
| **E) Linear regression** | Slope is zero (fit to means at each point) | 0.0468 |
| **Figure 5** |  |  |
| **C) Mann-Whitney U test** | Normalized mutual information is distributed the same if the temporal sequence of each cell's response is shuffled | <0.0001 |
| **G) Fisher's exact test** (groups numbered from left to right) | Proportion of stimuli eliciting an eruption is the same across genotypes | 1-2: 0.0002, 1-3: <0.0001, 2-3: 0.0784 |
| Mann-Whitney U test (groups numbered from left to right) | Eruption magnitude is the same across genotypes | 1-2: 0.016, 1-3: 0.0085, 2-3: 0.4136 |
| **Figure 6** |  |  |
| **A) Fisher's exact test** | Proportion long matings is the same across conditions | 30 sec relief: 1.0<br>90 sec relief: <0.0001 |
| **B) Fisher's exact test** | Proportion long matings is the same whether bPAC is activated or not | 1 min:<br>10 min: 0.0002 |
| **C) Fisher's exact test** | Proportion long matings is the same as the no light condition | 20 sec pulse: 0.111<br>40 sec pulse: <0.0001 |
| **Figure 7** |  |  |
| **D) Linear regressions** | Each regression is generated from the same linear trend | 0.7614 |
| **Figure S1** |  |  |
| **B) Correlation** | Decay time between simultaneously measured neurons is uncorrelated | 0.4670 |
| **B) Correlation** | Decay time between separately measured neurons is uncorrelated | 0.4683 |
| **Figure S4** |  |  |
| **B) Fisher's exact test** | Proportion terminating mating is no different between bPAC and Chrimson stimulation | <0.0001 |
| **Figure S5** |  |  |
| **D) Mann-Whitney U test** | Copulation durations are the same | <0.0001 |
| **Figure S6** |  |  |
| **C) Linear regression** | No slope in the relationship between pulse number and mean peak fluorescence change | 0.0274 |
| **D) Correlation (Pearson)** | Fluorescence lifetime change is not correlated with change in fluorescence | 0.0414 |
| **E) Correlation (Pearson)** | Fluorescence lifetime change is not correlated with change in fluorescence (using several cells in addition to exemplar trace) | 0.6593 |
| **Figure S7** |  |  |
| **B) F-statistic** | Slopes of all mean peak heights are the same across treatments | 0.014 |
| **D) Linear regression** | Slope of relationship between peak cAMP and pulse number is 0 | 0.0713 |
|  | Slope of relationship between residual cAMP and pulse number is 0 | 0.0335 |
|  | Slope of relationship between peak calcium and pulse number is 0 | .0689 |
|  | Slope of relationship between residual calcium and pulse number is 0 with residual calcium increase | 0.0087 |
| **Figure S8** |  |  |
| **C) Mann-Whitney U test** | Distribution of no light peak response is the same | WT – PKA-R*: >0.999<br>PKA-R* - CaMKII-T287D: 0.147<br>WT – CaMKII-T287D: **0.0173** |
|  | Distribution of 500 ms peak response is the same | WT – PKA-R*: **0.0109**<br>PKA-R* -- CaMKII-T287D: 0.317<br>WT – CaMKII-T287D: **0.0003** |
|  | Distribution of 1 sec peak response is the same | WT – PKA-R*: **0.0109**<br>PKA-R* -- CaMKII-T287D: 0.317<br>WT – CaMKII-T287D: **0.0003** |
|  | Distribution of 2 sec peak response is the same | WT – PKA-R*: **0.0097**<br>PKA-R* -- CaMKII-T287D: 0.359<br>WT – CaMKII-T287D: **0.0004** |
|  | Distribution of 5 sec peak response is the same | WT – PKA-R*: 0.0398<br>PKA-R* -- CaMKII-T287D: 0.191<br>WT – CaMKII-T287D: **0.0008** |
|  | Distribution of 7 sec peak response is the same | WT – PKA-R*: 0.0412<br>PKA-R* -- CaMKII-T287D: 0.288<br>WT – CaMKII-T287D: **0.0018** |
|  | Distribution of no light sustained response is the same | WT – PKA-R*: 0.0941<br>PKA-R* - CaMKII-T287D: 0.391<br>WT – CaMKII-T287D: 0.347 |
|  | Distribution of 500 ms sustained response is the same | WT – PKA-R*: 0.2415<br>PKA-R* -- CaMKII-T287D: 0.1760<br>WT – CaMKII-T287D: **0.015** |
|  | Distribution of 1 sec sustained response is the same | WT – PKA-R*: 0.1788<br>PKA-R* -- CaMKII-T287D: 0.1401<br>WT – CaMKII-T287D: **0.0065** |
|  | Distribution of 2 sec sustained response is the same | WT – PKA-R*: 0.488<br>PKA-R* -- CaMKII-T287D: 0.086<br>WT – CaMKII-T287D: 0.0233 |
|  | Distribution of 5 sec sustained response is the same | WT – PKA-R*: 0.6324<br>PKA-R* -- CaMKII-T287D: 0.0930<br>WT – CaMKII-T287D: 0.0438 |
|  | Distribution of 7 sec sustained response is the same | WT – PKA-R*: 0.3223<br>PKA-R* -- CaMKII-T287D: 0.1153<br>WT – CaMKII-T287D: **0.0144** |
| **Figure S10** |  |  |
| **A)** Mann-Whitney U test | Application of KN93 does not affect peak fluorescence change | CaMKII-T287D: **0.0055**<br>CaMKII-K43M-T287D: 0.8329 |
| **C)** Fisher's exact test | Probability of eliciting an eruption is the same with and without KN93 | **<0.0001** |

## Acknowledgements

We thank Andrew Siliciano and Vanessa Ruta for contribution of the unpublished cADDis reagents. We thank Yao Chen and Bernardo Sabatini for assistance with early microscopy experiments. We thank Mark Andermann and the members of the Crickmore and Rogulja labs for comments on the manuscript. We thank Eliza Smith, Jingwen Ren, and Isaac Lee for their assistance with the screen **Figure 2D**. The microscope used in these studies was constructed with the assistance of Ryohei Yasuda and Long Yan at Florida Lifetime Imaging. Funding:NIH R01GM134222 and an NSF graduate research fellowship to SCT.

## Author contributions

M.J.P. performed all behavioral experiments with assistance from S.C.T., C.S.A., and M.A.C. S.C.T. performed all microscopy experiments. Analysis and statistics code was written by S.C.T. The manuscript was written by S.C.T, M.J.P., and M.A.C. with input from C.A. All authors participated in analysis and interpretation of data.

